# Systemic muscle wasting and coordinated tumour response drive tumourigenesis

**DOI:** 10.1101/785022

**Authors:** Holly Newton, Yi-Fang Wang, Laura Camplese, André E.X. Brown, Susumu Hirabayashi

## Abstract

Cancer cells demand excess nutrients to support their proliferation, but how tumours exploit extracellular amino acids during systemic metabolic pertubations remain incompletely understood. Here we use a *Drosophila* model of obesity-enhanced tumourigenesis to uncover a systemic host-tumour nutrient circuit that supports tumour growth. We demonstrate coordinate induction of systemic cachexia-like muscle wasting with tumour-autonomous SLC36-family amino acid transporter expression as a proline-scavenging programme to drive tumourigenesis. This coordinated induction of cachexia and SLC36-transporters pertains to human kidney cancer and associates with significantly worse survival outcomes. We identify Indole-3-propionic acid as an optimal amino acid derivative to rationally target the proline-dependency of tumour growth. Combining insights from whole-animal *Drosophila* models and human cancer database analysis provides a powerful approach towards the identification and therapeutic exploitation of the amino acid vulnerabilities of tumourigenesis in the context of perturbed systemic metabolic network.

---

Cancer cells require a constant supply of metabolic intermediates to support their proliferation. To meet the biosynthetic demands associated with tumourigenesis, cancer cells actively acquire nutrients from the extracellular space^1-5^. Cancer is a systemic disease that associates with a range of host metabolic abnormalities such as obesity, insulin resistance, and cancer-associated cachexia; each of which alter the host systemic nutritional environment. These changes in both nutrient composition and availability may have profound effects on cancer development and progression. However, how cancer cells sense and respond to nutritional changes in the context of organismal metabolic alterations remains an under-explored area in cancer biology.

Cancer-associated cachexia is a systemic metabolic syndrome of weight loss associated with progressive skeletal muscle wasting^6,7^. The multifactorial and heterogeneous condition of cachexia involves a complex multi-organ interplay, which has impeded its comprehensive understanding at the molecular level^8^. It remains an intriguing open question in the field as to whether cancer-associated cachexia is an innocent bystander, or an active metabolic programme designed to meet the nutrient demands of tumourigenesis. Here we leverage a *Drosophila* model of obesity-enhanced tumourigenesis to directly address this question in the whole-animal setting. We demonstrate that obesity-enhanced tumours require proline for their growth, and that tumours induce systemic muscle wasting and SLC36-family transporter expression as a coordinated mechanism to exploit exogenous proline for tumourigenesis. We link these mechanistic insights from our fly study with human cancer data, to rationally target the proline-dependency of tumours as an approach to suppress tumour growth in the context of perturbed systemic metabolic network.

## Obesity-enhanced tumours induce muscle wasting

We previously reported a *Drosophila* larval model to study the systemic effects of diet-induced obesity and insulin resistance on tumour progression^9,10^. Feeding *Drosophila* larvae a high-sucrose diet (HSD) led to sugar-dependent metabolic defects including accumulation of fat, systemic insulin resistance, and hyperglycemia^11^. Targeted co-activation of Ras- and Src-pathways—by expression of an oncogenic isoform of *dRas1, ras1*^*G12V*^ and knockout of the negative regulator of Src, C-terminal src kinase (*csk*^*-/-*^)—in the *Drosophila* eye epithelia led to the development of benign tumours in animals raised on a control diet (CD) (Fig. 1a). Comparatively, feeding *ras1*^*G12V*^;*csk*^*-/-*^ animals a HSD promoted aggressive tumour growth in the eye epithelia (Fig. 1b), associated with secondary tumour formation (Fig. 1b, arrowheads), and led to larval lethality (Fig. 1c).

**Figure 1.**
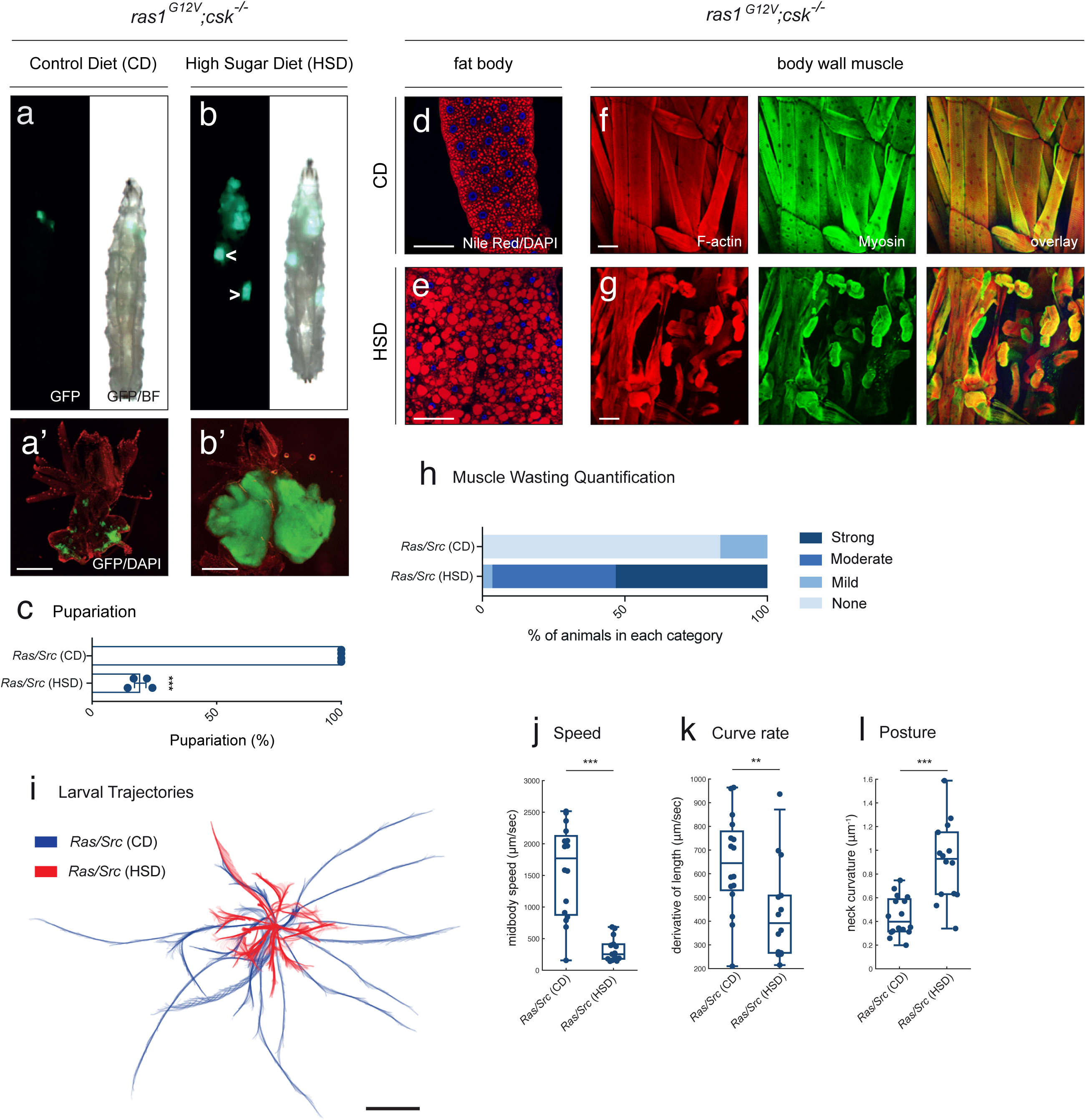
HSD-fed Ras/Src-animals exhibit cachexia-like muscle wasting and associated locomotion defects. **a, b**, *ras1*^*G12V*^;*csk*^*-/-*^ third-instar larvae fed a CD (**a**) or HSD (**b**). Transformed tissue is labelled with GFP (green). Secondary tumours are observed in a subset of animals (arrowheads in **b**). **a’, b’**, Matching dissected eye epithelial tissue stained with DAPI (red). Scale bar, 250 μm. **c**, Pupariation percentage of *ras1*^*G12V*^;*csk*^*-/-*^ animals raised on CD or HSD. Results are shown as mean ± SEM of four independent experiments with n≥10 animals for each data point. Asterisks indicate statistically significant difference (***; p<0.005). **d, e**, Nile Red (red) and DAPI (blue) staining of dissected fat body tissue from *ras1*^*G12V*^;*csk*^*-/-*^ third-instar larvae fed a CD (**d**) or HSD (**e**). Scale bar, 100 μm. **f, g**, F-actin (red) and anti-myosin (green) staining of dissected larval body wall muscle tissue from *ras1*^*G12V*^;*csk*^*-/-*^ third-instar larvae fed a CD (**f**) or HSD (**g**). Scale bar, 100 μm. **h**, Matching body wall muscle wasting quantification. n≥30 per condition. **i**, Individual larval trajectories from video tracking analysis of *ras1*^*G12V*^;*csk*^*-/-*^ third-instar larvae fed a CD, or HSD. Scale bar, 1 cm. **j-l**, Average midbody speed (**j**), body curvature (**k**), and body posture (**l**) of *ras1*^*G12V*^;*csk*^*-/-*^ third-instar larvae fed a CD, or HSD with results shown as mean ± SEM with each data point representing one individual animal. Asterisks indicate statistically significant difference (***; p<0.005, **; 0.005<p<0.025).

To explore the interplay between diet-induced obesity, tumour progression, and systemic organ wasting, we assessed the peripheral tissues of *ras1*^*G12V*^;*csk*^*-/-*^ animals—including the larval fat body and body wall skeletal muscle. Fat body lipid accumulation was elevated under HSD feeding, as assessed by the neutral lipid stain Nile Red (Fig. 1d, e). Strikingly, despite possessing an intact fat body, *ras1*^*G12V*^;*csk*^*-/-*^ animals raised on HSD exhibited significant skeletal muscle wasting as visualised by F-actin and myosin staining (Fig. 1f, g). To quantify the severity of muscle wasting—as assessed by imaging the fourth abdominal segment of the larval body wall muscle (Extended Data Fig. 1a)—we scored animals into one of four categories: none, minor, moderate or strong wasting (Extended Data Fig. 1b-e). While most (83.33%) of the *ras1*^*G12V*^;*csk*^*-*^ /^*-*^ animals raised on a CD displayed intact skeletal muscle, 100% of animals raised on a HSD exhibited muscle wasting of some degree (Fig. 1h). Muscle wasting was progressive; intact skeletal muscle was observed at the earlier stages of larval development, and as diet-enhanced tumourigenesis proceeded, skeletal muscle wasting increased (Extended Data Fig. 1f-j). Functional locomotion defects—as measured by video tracking analysis^12,13^—were observed in *ras1*^*G12V*^;*csk*^*-/-*^ animals raised on a HSD (Fig. 1i). Significant defects were observed in average speed, body curvature and body posture (Fig. 1j-l).

HSD feeding in *ras1*^*G12V*^;*csk*^*-/-*^ animals led to a developmental delay in reaching the third instar larval stage^9^. To rule out the possibility that observed muscle wasting was due to an extended larval period, we ablated the prothoracic gland (PG)—an ecdysteroid-producing organ—using the PG-specific *phantom (phm)-GAL4* driver to suppress pupariation. PG-ablated larvae, that were raised on a HSD and chronologically age-matched (day 6 after PG-ablation) to *ras1*^*G12V*^;*csk*^*-/-*^ animals in HSD, exhibited minimal muscle wasting in a HSD (Extended Data Fig. 2a-c). Together these results indicate that extended larval period alone is insufficient to account for the muscle wasting observed in *ras1*^*G12V*^;*csk*^*-/-*^ animals in HSD, and instead support a specific role for diet-enhanced tumourigenesis in promoting muscle wasting.

## Muscle wasting releases amino acids

A consequence of skeletal muscle wasting is protein degradation and subsequent release of free amino acids into the circulation. We performed a genome-wide transcriptional profiling analysis of dissected body-wall muscle from *ras1*^*G12V*^;*csk*^*-/-*^ animals raised on CD and HSD, from early through to late stages of muscle wasting (Extended Data Fig. 3a). As expected, Gene Set Enrichment Analysis (GSEA) revealed a progressive deregulation of muscle structure organisation in *ras1*^*G12V*^;*csk*^*-/-*^ animals in HSD (Extended Data Fig. 3b). A previous report demonstrated that ubiquitin-proteasome system (UPS) deregulation promotes skeletal protein degradation and muscle wasting via induction of the unfolded protein response (UPR)^14^. Consistent with this report, GSEA from muscle RNA-sequencing data revealed aberrant UPS function (Extended Data Fig. 3c) which correlated with accumulation of poly-ubiquitinated proteins (Extended Data Fig. 3d, e). This was coupled with an increase in UPR (Extended Data Fig. 3f), as visualised by increased perinuclear staining of the UPR-marker GRP-78^15^ (Extended Data Fig. 3g, h).

Through targeted metabolomic analysis, we measured an increase in circulating levels of amino acids in the hemolymph of *ras1*^*G12V*^;*csk*^*-/-*^ animals raised on a HSD relative to CD (Fig. 2a and Extended Data Fig. 4a, b). Together with increased circulating 3-methylhistidine, a biomarker for muscle atrophy^16^ (Extended Data Fig. 4c), these results indicate that muscle protein degradation takes place in *ras1*^*G12V*^;*csk*^*-/-*^ animals raised on a HSD. Since amino acids are one of the largest contributors to increased cell mass in proliferating cells^1^, our observations suggested muscle wasting as a source of circulating amino acids for tumour progression.

**Figure 2.**
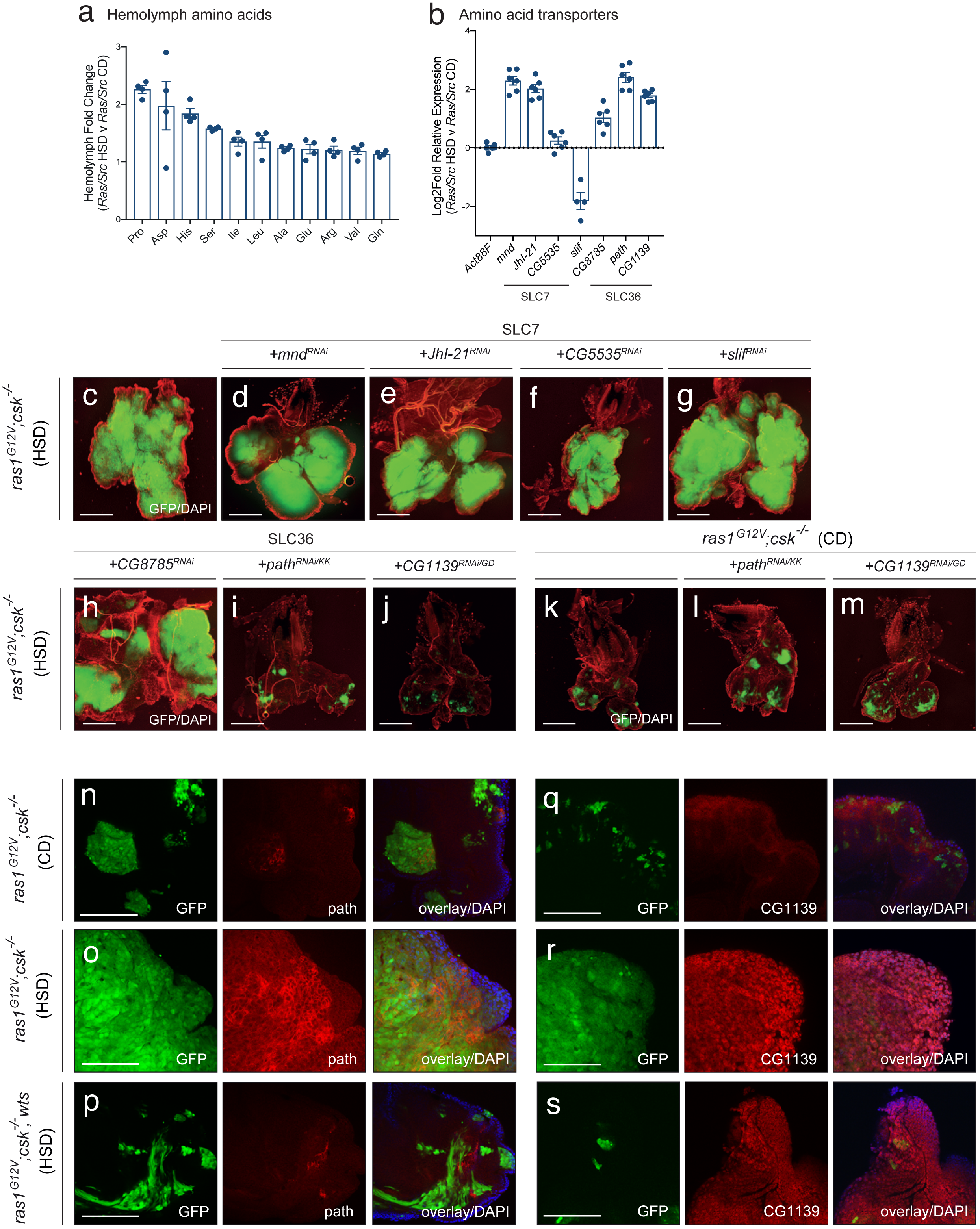
*Drosophila* SLC36-family amino acid transporters Path and CG1139 are required for obesity-enhanced Ras/Src tumour growth. **a**, Circulating hemolymph levels of upregulated amino acids in *ras1*^*G12V*^;*csk*^*-/-*^ animals raised on HSD compared to *ras1*^*G12V*^;*csk*^*-/-*^ animals raised on CD. Results are shown as mean relative fold change from *ras1*^*G12V*^;*csk*^*-/-*^ animals raised on HSD vs. CD. **b**, Relative log2fold change of SLC7 and SLC36 family amino acid transporters in dissected tumour tissue from *ras1*^*G12V*^;*csk*^*-/-*^ animals raised on HSD compared to *ras1*^*G12V*^;*csk*^*-/-*^ animals raised on CD, as determined by qPCR. Samples are normalised to *Act88F*. Results are shown as mean ± SEM. **c-j**, Dissected eye epithelial tissue stained with DAPI (red) from *ras1*^*G12V*^;*csk*^*-/-*^ animals raised on HSD (**c**), with *mnd*^*RNAi*^ (**d**), *JhI-21*^*RNAi*^ (**e**), *CG5535*^*RNAi*^ (**f**), *slif*^*RNAi*^ (**g**), *CG8785*^*RNAi*^ (**h**), *path*^*RNAi*^ (**i**), and *CG1139*^*RNAi*^ (**j**). Scale bar, 250 μm. **k-m**, Dissected eye epithelial tissue stained with DAPI (red) from *ras1*^*G12V*^;*csk*^*-/-*^ animals raised on CD (**k**), with *path*^*RNAi*^ (**l**), and *CG1139*^*RNAi*^ (**m**). Scale bar, 250 μm. **n-p**, Anti-Path staining (red) of dissected tumour tissue from *ras1*^*G12V*^;*csk*^*-/-*^ animals raised on CD (**n**), *ras1*^*G12V*^;*csk*^*-/-*^ animals raised on HSD (**o**), and *ras1*^*G12V*^;*csk*^*-/-*^,*wts* animals raised on HSD (**p**) with DAPI (blue). Scale bar, 100 μm. **q-s**, Anti-CG1139 staining (red) of dissected tumour tissue from *ras1*^*G12V*^;*csk*^*-/-*^animals raised on CD (**q**), *ras1*^*G12V*^;*csk*^*-/-*^ animals raised on HSD (**r**), and *ras1*^*G12V*^;*csk*^*-/-*^,*wts* animals raised on HSD (**s**) with DAPI (blue). Scale bar, 100 μm.

## Proline transporters are required for tumour growth

To explore candidates that could be responsible for tumour-specific amino acid utilisation, we focused on amino acid transporters. Through RNA-sequencing analysis of tumour tissue from *ras1*^*G12V*^;*csk*^*-/-*^ animals raised on CD and HSD, we characterised the expression profile of amino acid transporters across the SLC1, SLC7, SLC36 and SLC38 families (Extended Data Fig. 5a, 5b). Subsequent qPCR validation confirmed that amino acid transporters of the SLC7 family (*minidiscs (mnd), JhI-21* and CG5535) and SLC36-family (*CG8785, pathetic (path)*, and *CG1139*) were upregulated in tumours of *ras1*^*G12V*^;*csk*^*-/-*^ animals raised on a HSD compared to CD (Fig. 2b). Of note, expression levels of *Slimfast* (*slif*)—another SLC7-family member and an amino acid transporter previously implicated in tumour-specific usage of amino acids^17,18^—was downregulated in Ras/Src-tumours in HSD (Fig. 2b).

Reducing expression of *mnd (ras1*^*G12V*^;*csk*^*-/-*^,*mnd*^*RNAi*^*), JhI-21 (ras1*^*G12V*^;*csk*^*-/-*^,*JhI-21*^*RNAi*^*), CG5535 (ras1*^*G12V*^;*csk*^*-/-*^,*CG5535*^*RNAi*^*), slif (ras1*^*G12V*^;*csk*^*-/-*^,*slif*^*RNAi*^) or *CG8785* (*ras1*^*G12V*^;*csk*^*-/-*^, *CG8785*^*RNAi*^), had no effect on primary tumour size (Fig. 2c-h). Strikingly, reducing expression of *path* (*ras1*^*G12V*^,*path*^*RNAi*^;*csk*^*-/-*^) or CG1139 (*ras1*^*G12V*^;*csk*^*-/-*^,*CG1139*^*RNAi*^) almost completely suppressed diet-mediated Ras/Src-tumour growth (Fig. 2i, j and Extended Data Fig. 5c, d). Importantly, reduction in *path* or *CG1139* levels had minimal effect on tumour growth in *ras1*^*G12V*^;*csk*^*-/-*^ animals fed a CD (Fig. 2k-m) indicating that neither are required for benign tumour growth. Altogether, these results demonstrate that Path and CG1139 are specifically required for diet-mediated enhancement of Ras/Src-tumour growth.

We confirmed elevation of Path and CG1139 protein levels in Ras/Src-tumours in HSD using the Path- and CG1139-specific antibodies^19^ (Fig. 2n, o, q, r, and Extended Data Fig. 5e, f). *Path* has been previously identified as one of the growth-regulatory factors under control of the Hippo signalling pathway downstream transcriptional co-activator Yki^20^. Over-expression of Warts (Wts)—and therefore inhibition of Yki-activity—in Ras/Src-activated cells (*ras1*^*G12V*^;*csk*^*-/-*^,*wts*) led to decreased Path protein but not CG1139 levels in animals fed a HSD (Fig. 2p, s). While Path was elevated specifically in Ras/Src-activated tumours in HSD (Fig. 2n, o, and Extended Data Fig. 5e), the elevation of CG1139 was not specific to tumours but was rather dependent on high sugar diet (Fig. 2q, r, and Extended Data Fig. 5f). Together with our previous study demonstrating that diet-enhanced Ras/Src-tumours activate Yki^10^, we conclude that the Hippo signalling downstream effector Yki mediates *path* expression in *ras1*^*G12V*^;*csk*^*-/-*^ animals raised on a HSD.

## Proline promotes tumour growth via Path

Mammalian SLC36-family amino acid transporters including SLC36A1 and SLC36A2 transport alanine, glycine and proline^21^. Proline was the most highly increased amino acid in the hemolymph of *ras1*^*G12V*^;*csk*^*-/-*^ animals in HSD (Fig. 2a and Extended Data Fig. 4b). Over-expression of Path (*ras1*^*G12V*^;*csk*^*-/-*^,*pathA*) or CG1139 (*ras1*^*G12V*^,*CG1139;csk*^*-/-*^) had a negligible effect on tumour growth in Ras/Src-activated cells on CD, indicating that transporter expression alone is insufficient to promote tumourigenesis (Fig. 3a-c).

**Figure 3.**
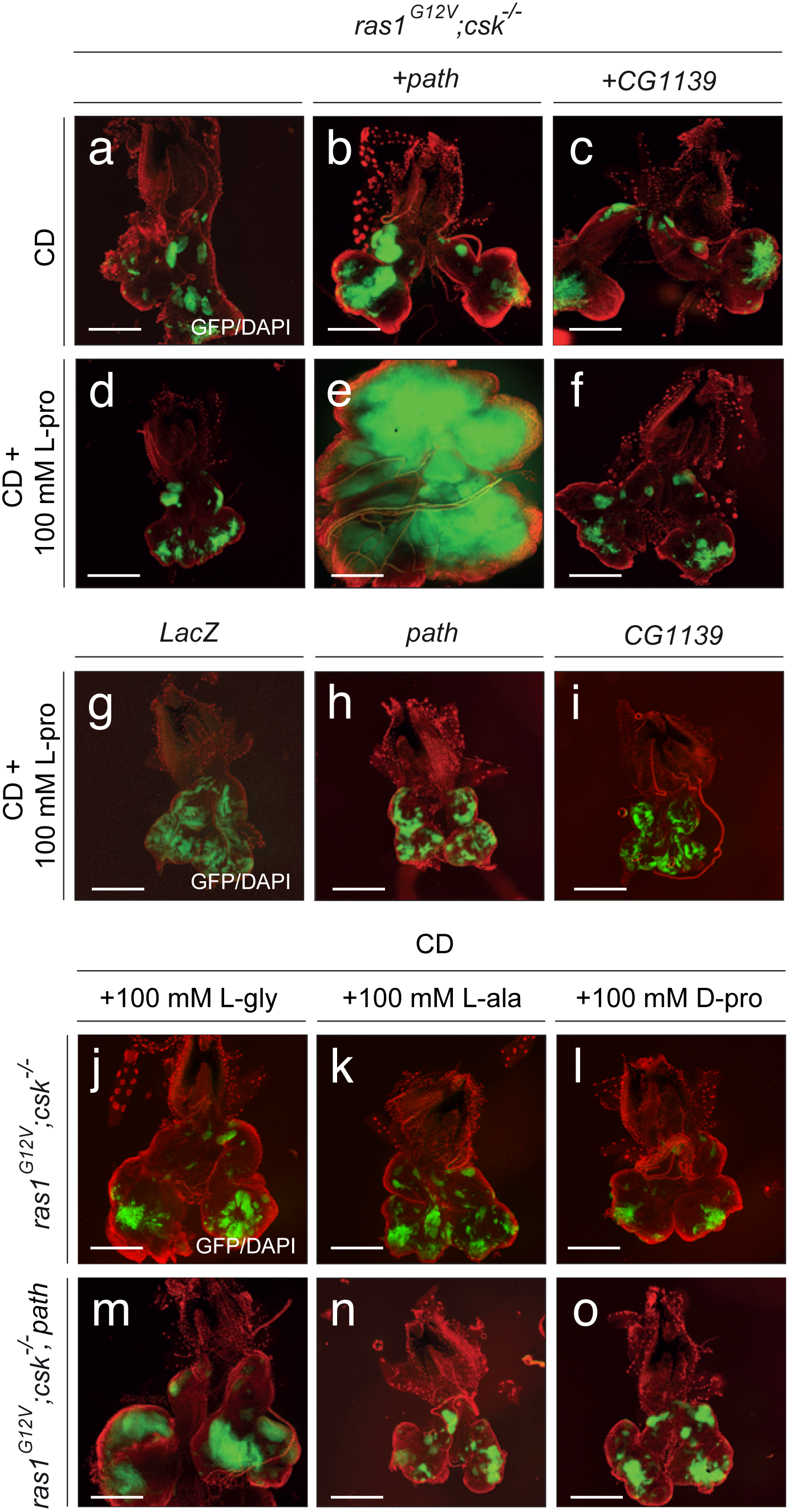
Proline promotes Ras/Src-tumour growth through amino acid transporter Path. **a-c**, Dissected eye epithelial tissue stained with DAPI (red) from *ras1*^*G12V*^;*csk*^*-/-*^ animals (**a**), *ras1*^*G12V*^;*csk*^*-/-*^,*PathA* animals (**b**), and *ras1*^*G12V*^,*CG1139;csk*^*-/-*^ animals (**c**) raised on CD. Scale bar, 250 μm. **d-f**, Dissected eye epithelial tissue stained with DAPI (red) from *ras1*^*G12V*^;*csk*^*-/-*^animals (**d**), *ras1*^*G12V*^;*csk*^*-/-*^,*PathA* animals (day 10 AEL) (**e**), and *ras1*^*G12V*^,*CG1139;csk*^*-/-*^ animals (**f**) raised on CD supplemented with 100 mM L-proline. Scale bar, 250 μm. **g-i**, Dissected eye epithelial tissue stained with DAPI (red) from *LacZ* animals (**g**), *PathA* animals (**h**), and *CG1139* animals (**i**) raised on CD supplemented with 100 mM L-proline. Scale bar, 250 μm. **j-l**, Dissected eye epithelial tissue stained with DAPI (red) from *ras1*^*G12V*^;*csk*^*-/-*^ animals raised on CD supplemented with 100 mM L-glycine (**j**), 100 mM L-alanine (**k**), and 100 mM D-proline (**l**). Scale bar, 250 μm. **m-o**, Dissected eye epithelial tissue stained with DAPI (red) from *ras1*^*G12V*^;*csk*^*-/-*^,*PathA* animals raised on CD supplemented with 100 mM L-glycine (**m**), 100 mM L-alanine (**n**), and 100 mM D-proline (**o**). Scale bar, 250 μm.

To mimic the elevation of circulating proline levels under conditions of muscle wasting, we carried out dietary supplementation of proline. Feeding animals CD supplemented with 100 mM L-proline had minimal effect on tumour growth of *ras1*^*G12V*^;*csk*^*-/-*^ animals in CD (Fig. 3d). Intriguingly however, feeding proline in *ras1*^*G12V*^;*csk*^*-/-*^,*pathA* animals strongly promoted tumour growth in CD (Fig. 3e). In striking contrast, proline feeding in *ras1*^*G12V*^,*CG1139;csk*^*-/-*^ animals had no effect on tumour growth (Fig. 3f). Of note, feeding proline in animals with path- or CG1139-overexpressing clones had minimal effect on tissue growth (Fig. 3g-i), suggesting that proline-Path mediated enhancement of tissue growth requires an oncogenic background. Altogether, these results indicate that coordinated elevation of systemic proline availability and tumour-autonomous transporter expression is required to promote Ras/Src-tumour growth.

Feeding excess L-glycine or L-alanine had minimal growth promoting effect in *ras1*^*G12V*^;*csk*^*-/-*^ and *ras1*^*G12V*^;*csk*^*-/-*^,*pathA* animals (Fig. 3j, k, m, n). Similarly, feeding the biologically inert isomer of proline—D-proline—had no effect on tumour growth in both *ras1*^*G12V*^;*csk*^*-/-*^ and *ras1*^*G12V*^;*csk*^*-/-*^,*pathA* animals (Fig. 3l, o), thereby indicating that tumour response is specific to L-proline. Altogether, our data highlight the proline vulnerability of obesity-enhanced Ras/Src-tumours and uncover modulation of amino acid transporter repertoire as a strategy to meet the nutrient requirements of tumourigenesis.

## Muscle wasting affects tumour growth

Our results suggest that cachexia-like muscle wasting and associated circulating proline availability promote Ras/Src-tumourigenesis. Therefore, targeting tumour-derived factors which mediate muscle wasting may not only attenuate muscle wasting but also suppress tumour growth. To explore this concept, we set out to identify tumour-derived factors which mediate muscle wasting. Through genome-wide transcriptional profiling analysis of dissected tumour tissue from *ras1*^*G12V*^;*csk*^*-/-*^ animals raised on CD and HSD (Extended Data Fig. 5a), we identified *branchless* (*bnl*), a *Drosophila* fibroblast growth factor (FGF) ligand as one of the most highly upregulated secreted factors in *ras1*^*G12V*^;*csk*^*-/-*^ tumours in animals raised on a HSD (upregulated 5.02-log2fold)(Fig. 4a and Extended Data Fig. 6a). Among the three known *Drosophila* FGF ligands (Bnl, Pyramus, and Thisbe), *bnl* was the only FGF ligand to be highly elevated in *ras1*^*G12V*^;*csk*^*-/-*^ tumours in HSD (Fig. 4a). Consistently, Bnl protein expression was strongly upregulated in *ras1*^*G12V*^;*csk*^*-/-*^ tumours of animals fed a HSD (Fig. 4b, c).

**Figure 4.**
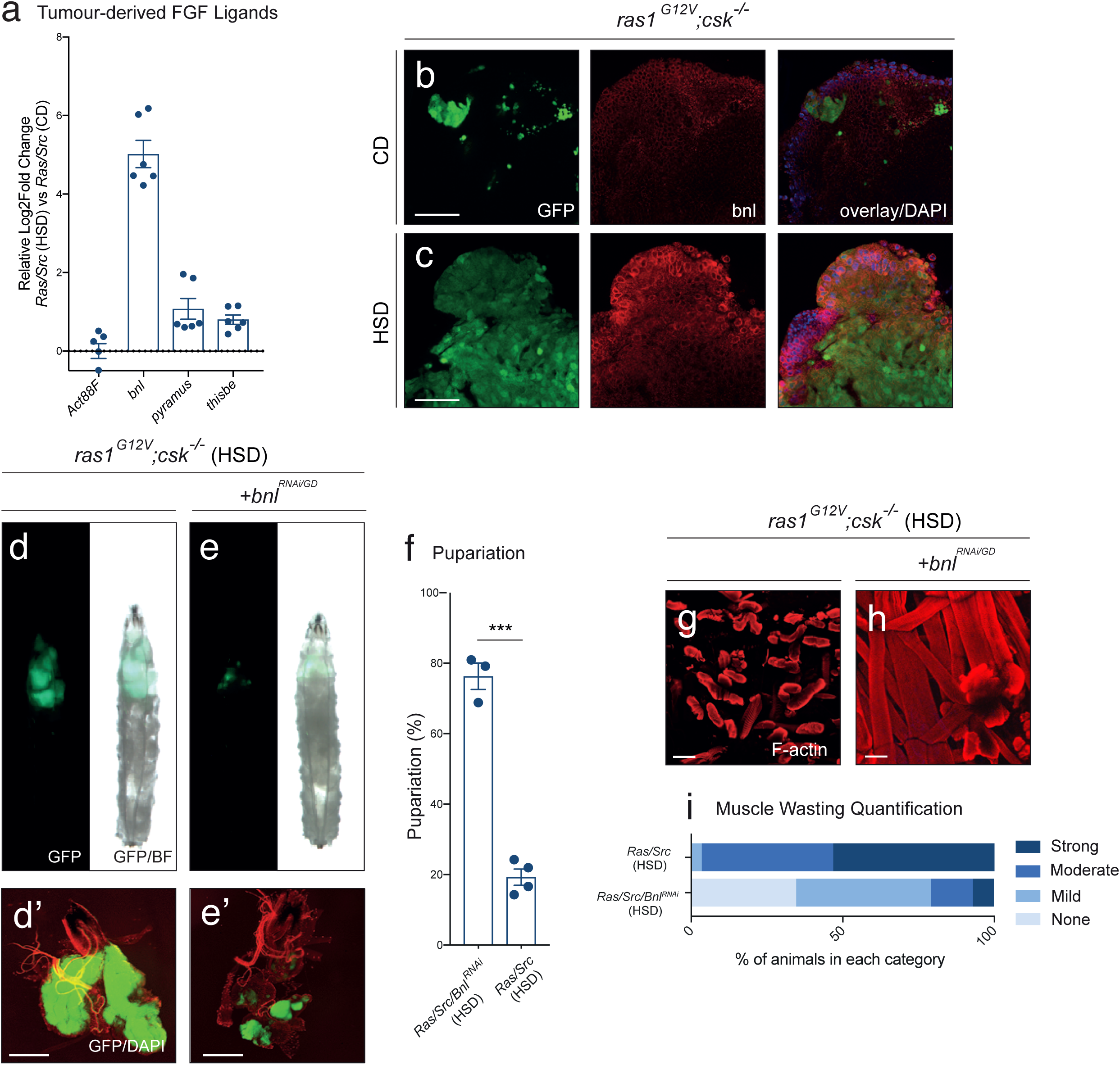
Tumour-derived branchless mediates muscle wasting and subsequent tumour growth. **a**, Relative log2fold change of *Drosophila* fibroblast growth factors, *branchless* (*bnl*), *pyramus* and *thisbe* in dissected tumour tissue from *ras1*^*G12V*^;*csk*^*-/-*^ animals raised on HSD compared to *ras1*^*G12V*^;*csk*^*-/-*^ animals raised on CD, as determined by qPCR. Samples are normalised to *Act88F*. Results are shown as mean ± SEM. **b, c**, Anti-Bnl staining (red) of dissected tumour tissue from *ras1*^*G12V*^;*csk*^*-/-*^ animals raised on CD (**b**) or HSD (**c**) with DAPI (blue). Scale bar, 40 μm. **d, e**, Third-instar larvae from *ras1*^*G12V*^;*csk*^*-/-*^ (**d**), and *ras1*^*G12V*^;*csk*^*-/-*^,*bnl*^*RNAi*^ (**e**). **d’, e’**, Matching dissected eye epithelial tissue stained with DAPI (red). Scale bar, 250 μm. **f**, Pupariation percentage of animals raised on HSD. Results are shown as mean ± SEM of independent experiments with n≥10 animals for each data point. Asterisks indicate statistically significant difference (***; p<0.005). **g, h**, Matching dissected larval body wall muscle. Scale bar, 100 μm. **i**, Matching body wall muscle wasting quantification. n≥30 per condition.

Reducing *bnl* expression in Ras/Src-activated tumours (*ras1*^*G12V*^;*csk*^*-/-*^,*bnl*^*RNAi*^) attenuated muscle wasting in animals raised on a HSD (Fig. 4g-i and Extended Data Fig. 6f, k), indicating that tumour-derived Bnl is required for muscle wasting. Tumour-derived factors recently implicated in cachexia-like organ wasting include ImpL2 and Pvf1^22-24^. These were also elevated both transcriptionally and at the protein level in *ras1*^*G12V*^;*csk*^*-/-*^ tumours in HSD (Extended Data Fig. 6a-e). However, reducing the levels of either ImpL2 (*ras1*^*G12V*^;*csk*^*-/-*^,*ImpL2*^*RNAi*^) or Pvf1 (*ras1*^*G12V*^,*pvf1*^*RNAi*^;*csk*^*-/-*^) failed to rescue muscle wasting as efficiently as *bnl* in our diet-enhanced Ras/Src-tumour model (Extended Data Fig. 6g, h, k). Altogether, these results establish Bnl as a functionally relevant tumour-secreted factor that contributes to muscle wasting in animals with diet-enhanced Ras/Src-tumours.

Importantly, reducing *bnl* expression in diet-enhanced Ras/Src-tumours (*ras1*^*G12V*^;*csk*^*-/-*^,*bnl*^*RNAi*^) not only attenuated systemic muscle wasting but also suppressed tumour growth and larval lethality (Fig. 4d-f and Extended Data Fig. 6f). Tumour autonomous Bnl and its receptor Breathless (Btl) signalling has been implicated in tumourigenesis^25^. However, reducing the levels of *btl* in Ras/Src-activated tumours (*ras1*^*G12V*^,*btl*^*RNAi*^;*csk*^*-/-*^) had minimal effect on primary tumour size or muscle wasting, indicating that tumour autonomous reduction of Bnl-Btl signalling was not responsible for suppression of tumour growth in *ras1*^*G12V*^;*csk*^*-/-*^,*bnl*^*RNAi*^ animals (Extended Data Fig. 6i-k). Together, these results reveal that systemic muscle wasting has a functional effect on tumour growth.

We next examined the link between proline-mediated Ras/Src tumour growth and Bnl-mediated muscle wasting. Feeding proline in *ras1*^*G12V*^;*csk*^*-/-*^,*pathA* animals not only promoted tumour growth (Fig. 3e), but also led to elevated tumour expression of Bnl (Extended Data Fig. 7a, b) and subsequent muscle wasting (Extended Data Fig. 7c-e). Together this demonstrates that Path-dependent proline-enhancement of tumour growth promotes Bnl-mediated muscle wasting. Overall, this data implicates the coordinate induction of amino acid transporter expression, coupled with cachexia-like muscle wasting, as sufficient to establish a feed-forward host-tumour circuit to drive tumour growth.

## Translation to human cancer

To examine whether coordinated induction of cachexia and amino acid transporter expression pertains to human cancers, we examined the transcriptional profile of tumour samples from patients in The Cancer Genome Atlas (TCGA), across a panel of cancer types^26^. As with most databases, patient data is not annotated for the presence or absence of cachexia, however an earlier study adopted the expression of cachectic factors—IL-1A, IL-1B, IL-6, IL-8 and TNF-α—as a proxy for cachexia (Cachexia Score)^27^. In humans, the SLC36 family of amino acid transporters consists of four members: SLC36A1-A4. We focused our analysis on SLC36A1 and SLC36A4; the A2 and A3 isoforms were excluded due to their low tumour expression levels. From the six cancer types analysed, Cachexia Score correlated positively with both SLC36A1 and SLC36A4 expression in kidney cancer, SLC36A1 expression in pancreatic cancer, and SLC36A4 expression in liver cancer patients (Fig. 5a-e). To determine whether coordinated induction of cachectic factors and SLC36-family transporters affected prognosis, we compared survival outcomes in patient groups with high and low expression of cachectic factors and SLC36 transporters. Strikingly, kidney cancer patients with both a high Cachexia Score and high SLC36 transporter expression had significantly worse survival compared with those exhibiting both a low Cachexia Score and low SLC36 transporter expression (Fig. 5f, g). Similarly patients with both a high Cachexia Score and a high SLC36A1 expression in pancreatic cancer or a high Cachexia Score and a high SLC36A4 expression in liver cancer exhibited worse survival rates, albeit not reaching statistical significance (Fig. 5h, i). Overall, our data demonstrates that coordinated induction of cachectic factors and SLC36 transporter expression has a strong negative effect on patient survival in human kidney cancer.

**Figure 5.**
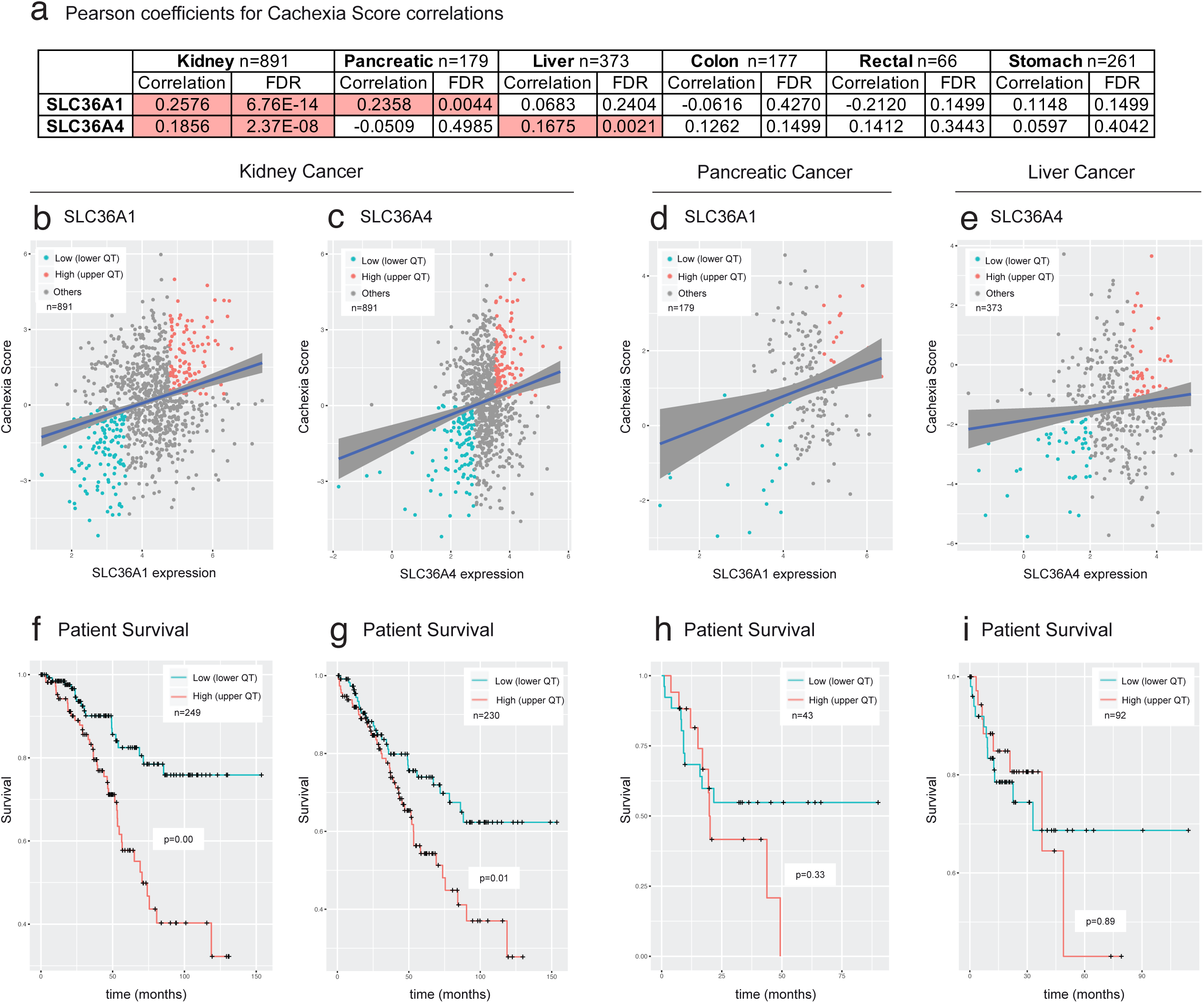
Coordinated induction of cachectic factors and SLC36-transporter expression decreases survival in kidney cancer patients. **a**, Pearson correlation coefficients and associated false discovery rates (FDR) for observed correlation between Cachexia Score and SLC36A1 or SLC36A4 expression levels across kidney, pancreatic, liver, colon, rectal, and stomach cancer patient groups. Red denotes conditions with statistically significant positive correlation (FDR<0.05). **b-e**, Scatter plot and overlaid linear regression curve for Cachexia Score and SLC36A1 (**b**) or SLC36A4 (**c**) expression levels in kidney cancer (n=891), Cachexia Score and SLC36A1 expression in pancreatic cancer (n=179) (**d**), and Cachexia Score and SLC36A4 expression in liver cancer (n=373) (**e**). Blue data points denote the lower quartile of patients with low Cachexia Score and low SLC36; red data points denote the upper quartile of patients with high Cachexia Score and high SLC36 transporter expression; grey data points denote the 2^nd^ and 3^rd^ quartile of patients classified as ‘Others’. **f-i**, Kaplan-Meier survival curve for the upper quartile of patients with high Cachexia Score and high transporter expression (High) and lower quartile of patients with low Cachexia Score and low transporter expression (Low). SLC36A1 patient groups in kidney cancer as defined in **b** (n=249) (**f**), SLC36A4 patient groups in kidney cancer as defined in **c** (n=230) (**g**), SLC36A1 patient groups in pancreatic cancer as defined in **d** (n=43) (**h**), and SLC36A4 patient groups in liver cancer as defined in **e** (n=92) (**i**).

## Targeting proline vulnerability of tumours

Taken together, our *Drosophila* data and human cancer database analysis specifically highlight SLC36-family transporters as a target for therapeutic intervention in tumours with proline vulnerability. A previous *in vitro* study using a human cancer cell line explored amino acid- and amino acid derivative-specificities of SLC36A1^28^. By selecting competitive, non-transported inhibitors of SLC36A1 from this study, we performed a whole-animal *Drosophila* screen to identify candidates that suppress tumour growth and display minimal whole-animal toxicity. Feeding *ras1*^*G12V*^;*csk*^*-/-*^ animals a HSD supplemented with 5-Hydroxy-L-tryptophan led to animal lethality at the early larval stages, indicative of whole-animal toxicity. Feeding *ras1*^*G12V*^;*csk*^*-/-*^ animals a HSD supplemented with either L-tryptophan or Indole-5-carboxylic acid led to successful larval development, but had no inhibitory effect on tumour growth (Figure 6a-c). However, feeding *ras1*^*G12V*^;*csk*^*-/-*^ animals a HSD supplemented with Indole-3-propionic acid (IPA) dramatically suppressed tumour growth in HSD (Fig. 6d). Strikingly, 66.2% of IPA-fed *ras1*^*G12V*^;*csk*^*-/-*^ animals achieved pupariation in HSD, revealing significant tumour-suppressing efficacy coupled with minimal whole-animal toxicity (Fig. 6g-i). Furthermore, IPA feeding had no effect on benign tumour growth in *ras1*^*G12V*^;*csk*^*-/-*^ animals raised on CD (Fig. 6e, f), supporting the notion that SLC36 inhibition only limits the growth of tumours with proline vulnerability.

**Figure 6.**
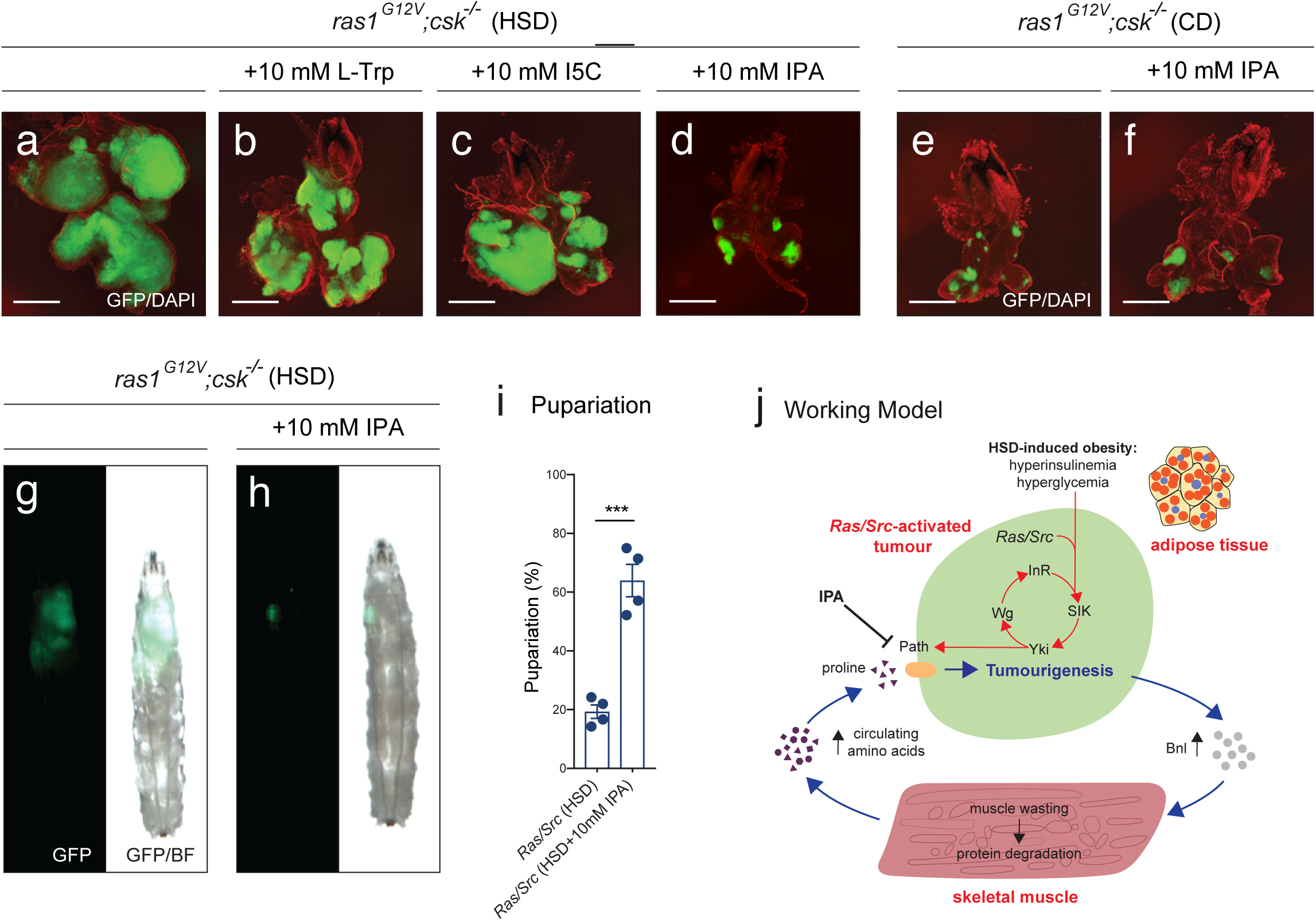
Targeting proline vulnerability with amino acid derivatives suppresses tumour growth. **a-d**, Dissected eye epithelial tissue stained with DAPI (red) from *ras1*^*G12V*^;*csk*^*-/-*^ animals raised on HSD (**a**), HSD supplemented with 10 mM L-tryptophan (**b**), HSD supplemented with 10 mM Indole-5-carboxylic acid (**c**), HSD supplemented with 10 mM Indole-3-propionic acid (IPA) (**d**). Scale bar, 250 μm. **e, f**, Dissected eye epithelial tissue stained with DAPI (red) from *ras1*^*G12V*^;*csk*^*-/-*^ animals raised on CD (**e**), and CD supplemented with 10 mM IPA (**f**). Scale bar, 250 μm. **g, h**, Third-instar larvae from *ras1*^*G12V*^;*csk*^*-/-*^ animals raised on HSD (**g**) and HSD supplemented with 10 mM Indole-3-propionic acid (IPA) (**h**). **i**, Pupariation percentage of *ras1*^*G12V*^;*csk*^*-/-*^ animals raised on HSD and HSD supplemented with 10 mM IPA. Results are shown as mean ± SEM of four independent experiments with n≥10 animals for each data point. Asterisks indicate statistically significant difference (***; p<0.005). **j**, Working model illustrating dual-layered-coordination of tumour metabolic response: (i) at the whole organism level—Branchless (Bnl)-dependent induction of cachexia-like muscle wasting and systemic amino acid release (blue arrows)—and (ii) at the tumour-autonomous level—Yorkie-dependent induction of amino acid transporter expression (red arrows). The tumour-autonomous SIK-Yki-Wg-InR circuit was identified previously^10^. IPA represents a therapeutic strategy to break the nutritional circuit between muscle wasting and tumour growth by targeting the SLC36 family transporter Path. SIK; Salt-inducible kinase, Yki; Yorkie, Wg; Wingless, InR; Insulin Receptor, IPA; indole-3-propionic acid.

## DISCUSSION

Our study reveals a systemic amino acid-utilising circuit whereby obesity-enhanced tumours induce muscle wasting as a systemic metabolic network to drive tumourigenesis. A consequence of muscle wasting in *ras1*^*G12V*^;*csk*^*-/-*^ animals raised on HSD was increased release of proline into the circulation (Fig. 2a). Plasma amino acid profiling of patients with sarcopenia—a condition of muscle wasting associated with aging—showed elevated plasma proline levels^29^, indicating that elevated free circulating proline is a common feature of muscle wasting. We identify a proline vulnerability of obesity-enhanced tumours; SLC36-family proline transporter Path is required for tumour growth and exogenous proline promotes tumourigenesis through Path (Fig. 2, 3). We highlight two layers of coordination in tumour metabolic response: (i) at the whole organism level - by promoting cachexia-like muscle wasting and systemic amino acid availability, and (ii) at the tumour-autonomous level - by altering amino acid transporter repertoire (Fig. 6j).

The insights obtained from our *Drosophila* model were sufficient to predict altered cancer patient outcomes. Human cancer database analysis demonstrates that co-expression of cachectic factors alongside both SLC36A1 and SLC36A4 amino acid transporters correlates with significantly worse patient survival in kidney cancer patients (Fig. 5). Proline has been shown to be a limiting amino acid for protein synthesis in kidney cancers^30^. Therefore, reducing proline uptake with SLC36-inhibitors—as exemplified here by use of Indole-3-propionic acid (Fig. 6)— may warrant consideration as a therapeutic strategy to break the nutritional circuit between cachexia-like muscle wasting and tumour growth in human kidney cancer patients.

## METHODS

### Fly stocks

*UAS-ras1*^*G12V*^, *UAS-lacZ, UAS-bnl*^*RNAi/TRiP*^, *UAS-path*^*RNAi/TRiP*^ and *UAS-wts* flies were obtained from the Bloomington *Drosophila* Stock Center. *UAS-bnl*^*RNAi/GD*^, *UAS-path*^*RNAi/KK*^, *UAS-Pvf1*^*RNAi/GD*^, *UAS-mnd*^*RNAi/GD*^, *UAS-slif*^*RNAi/GD*^, *UAS-JhI-21*^*RNAi/GD*^, *UAS-ImpL2*^*RNAi/GD*^, *UAS-CG8785*^*RNAi/GD*^, *UAS-CG5535*^*RNAi/GD*^, *UAS-CG1139*^*RNAi/GD*^, *UAS-CG1139*^*RNAi/KK*^, *UAS-btl*^*RNAi/GD*^ and *UAS-btl*^*RNAi/KK*^ flies were obtained from the Vienna *Drosophila* Resource Center.

The following stocks were kindly provided to us: *FRT82B, csk*^*Q156Stop*^ by A. O’Reilly and M. Simon; *ey(3.5)-FLP1* by G. Halder; *Phm-gal4; tub-gal80*^*ts*^ and *UAS-grim* by T. Koyama; *UAS-pathA/TM6B* by J. Parrish; *UAS-CG1139* by D. Goberdhan.

To create eyeless-driven GFP-labeled clones, the MARCM system was used. Flies with the genotype *ey(3.5)-FLP1; act > y+ > gal4,UAS-GFP; FRT82B, tub-gal80* were crossed to flies with the following genotypes: (1) *UAS-ras1*^*G12V*^; *FRT82B, csk*^*Q156Stop*^*/TM6b*; (2) *UAS-ras1*^*G12V*^; *FRT82B, csk*^*Q156Stop*^, *UAS-mnd*^*RNAi/GD*^*/TM6b*; (3) *UAS-ras1*^*G12V*^; *FRT82B, csk*^*Q156Stop*^, *UAS-JhI-21*^*RNAi/GD*^*/TM6b*; (4) *UAS-ras1*^*G12V*^; *FRT82B, csk*^*Q156Stop*^, *UAS-CG5535*^*RNAi/GD*^*/TM6b*; (5) *UAS-ras1*^*G12V*^; *FRT82B, csk*^*Q156Stop*^, *UAS-slif*^*RNAi/GD*^*/TM6b*; (6) *UAS-ras1*^*G12V*^; *FRT82B, csk*^*Q156Stop*^, *UAS-CG8785*^*RNAi/GD*^*/TM6b*; (7) *UAS-ras1*^*G12V*^, *UAS-path*^*RNAi/KK*^; *FRT82B, csk*^*Q156Stop*^*/TM6b*; (8) *UAS-ras1*^*G12V*^; *FRT82B, csk*^*Q156Stop*^, *UAS-CG1139*^*RNAi/GD*^*/TM6b*; (9) *UAS-ras1*^*G12V*^, *UAS-path*^*RNAi/TRiP*^; *FRT82B, csk*^*Q156Stop*^*/TM6b*; (10) *UAS-ras1*^*G12V*^, *CG1139*^*RNAi/KK*^; *FRT82B, csk*^*Q156Stop*^*/TM6b*; (11) *UAS-ras1*^*G12V*^; *FRT82B, csk*^*Q156Stop*^, *UAS-wts*; (12) *UAS-ras1*^*G12V*^; *FRT82B, csk*^*Q156Stop*^, *UAS-PathA/TM6b*; (13) *UAS-CG1139, UAS-ras1*^*G12V*^; *FRT82B, csk*^*Q156Stop*^*/TM6b*; (14) *UAS-lacZ; FRT82B*; (15) *FRT82B, UAS-PathA/TM6b*; (16) *UAS-CG1139; FRT82B*; (17) *UAS-ras1*^*G12V*^; *FRT82B, csk*^*Q156Stop*^, *UAS-bnl*^*RNAi/GD*^*/SM6-TM6b*; (18) *UAS-ras1*^*G12V*^; *FRT82B, csk*^*Q156Stop*^, *UAS-bnl*^*RNAi/TRiP*^*/TM6b*; (19) *UAS-ras1*^*G12V*^, *UAS-btl*^*RNAi/GD*^; *FRT82B, csk*^*Q156Stop*^*/TM6b*; (20) *UAS-ras1*^*G12V*^, *UAS-btl*^*RNAi/KK*^; *FRT82B, csk*^*Q156Stop*^*/TM6b*; (21) *UAS-ras1*^*G12V*^; *FRT82B, csk*^*Q156Stop*^, *UAS-ImpL2*^*RNAi/GD*^*/TM6b*; (22) *UAS-ras1*^*G12V*^; *FRT82B, csk*^*Q156Stop*^, *UAS-Pvf1*^*RNAi/GD*^*/TM6b*.

To carry out prothoracic gland ablation, *Phm-gal4; tub-gal80*^*ts*^*/TM6b* flies were crossed with *UAS-grim* flies and reared at 18°C. At early third instar larval stage (day 12 in 18°C), cultures were switched to 29°C to inactivate *tub-gal80*^*ts*^.

### Cultures

Cultures were carried out on Bloomington semi-defined medium (https://bdsc.indiana.edu/information/recipes/germanfood.html) and contained sucrose (0.15 M and 1.0 M in CD and HSD, respectively) as the only purified sugar source. Ingredients were obtained from; Agar (Fisher Scientific; BP2641-1), Brewer’s Yeast (MP Biomedicals; 903312, Lot: BCBN0171V), Yeast Extract (Sigma-Aldrich; 70161), Peptone (Sigma-Aldrich; 82303), Sucrose (Fisher Scientific; S/8560/63), Magnesium sulfate hexahydrate (Fluka; 00627), Calcium chloride dihydrate (Sigma-Aldrich; 223506), Propionic acid (Sigma-Aldrich; P1386), p-Hydroxy-benzoic acid methyl ester (Sigma-Aldrich; H5501). For dietary supplementation experiments, L-proline (Sigma-Aldrich; P5607), D-proline (Sigma-Aldrich; 858919), L-glycine (VWR Chemicals; 101196X) or L-alanine (Sigma-Aldrich; 7627) were added to CD to a final concentration of 100 mM. L-tryptophan (Sigma Aldrich; T0254), Indole-5-carboxylic acid (Sigma Aldrich; I5400) and Indole-3-propionic acid (Sigma-Aldrich; 57400) were added to HSD and CD to a final concentration of 10 mM. Cultures were performed at 25°C unless otherwise noted. HSD feeding led to a developmental delay in reaching the third instar larval stage; all experiments were performed on late third instar larvae at day 8 after egg laying (CD) and day 14 after egg laying (HSD) unless otherwise noted.

### Immunohistochemistry

Tissue was dissected in PBS, fixed in 4% PFA/PBS for 30’ on ice, washed in PBS-T, incubated with primary antibody in PAXDG (PBS containing 1% BSA, 0.3% Triton X-100, 0.3% deoxycholate, and 5% goat serum), followed by washing and incubation with secondary antibody in PAXDG and subsequent mounting in Vectashield with DAPI (Vector Laboratories). Primary antibodies used were: rat anti-myosin (ab51098; Abcam; 1:100), mouse anti-polyubiquitin (FK2; Enzo Life Sciences; 1:300), mouse anti-GRP-78 (E-4; Santa Cruz Biotechnology; 1:50). Polyclonal Rabbit anti-CG1139 (1:200) was purified by New England Peptide Inc., MA, USA. An amino acid sequence 26-40 was selected as the epitope. The following primary antibodies were kindly gifted to us: rat anti-bnl (M. Krasnow; 1:50) guinea pig anti-path (J. Parrish; 1:200), rat anti-ImpL2 (H. Stocker; 1:250), rat anti-pvf1 (B. Shilo; 1:200). Secondary antibodies used were: Alexa-488 and Alexa-568 conjugated anti-rat antibody (Molecular Probes; 1:200), Alexa-488 and Alexa-568 conjugated anti-mouse antibody (Molecular Probes; 1:200), Alexa-488 and Alexa-568 conjugated anti-rabbit antibody (Molecular Probes; 1:200) and Alexa-568 conjugated anti-guinea pig antibody (Molecular Probes; 1:200). TRITC-labelled rhodamine-phalloidin was used to visualise F-actin (R415; Invitrogen; 1:500).

### Nile red staining

Tissue was dissected in PBS, fixed in 4% PFA/PBS for 30’ on ice, incubated with 10 μg/mL Nile Red stain (N1142; ThermoFisher Scientific) in PBS for 30’ on ice followed by washing and subsequent mounting in Vectashield with DAPI.

### Muscle wasting quantification

Individual larvae were washed and dissected in PBS. Larval body wall muscle was stained with TRITC-labelled rhodamine-phalloidin (R415; Invitrogen; 1:500) to visualise F-actin. Individual animals were scored into categories each representing the strength of muscle wasting from either “none”; “minor”; “moderate” or “strong” (Extended Data Fig. 1a-e). Scores were assigned based on wasting of the ventral longitudinal, lateral oblique and lateral longitudinal muscles centered around the 4^th^ abdominal segment (A4) of the larval body wall (Extended Data Fig. 1a). Each hemisegment was used as an internal control. A minimum of 30 animals were scored per condition. Body wall muscle staining and quantification was performed on animals from day 8 (CD), day 10 (early stage; HSD), day 14 (mid stage; HSD), and day 16 (late stage; HSD) after egg laying.

### Microscopy and imaging

Larval images were acquired with Leica M165 FC fluorescent microscope equipped with Leica DFC 3000G digital camera. Confocal images were taken with Leica SP5 II confocal microscope and Leica Application Suite Advanced Fluorescence software (Leica Microsystems). Larval GFP/brightfield overlay images were compiled with Adobe Photoshop software.

### Larval video tracking and analysis

Larvae were maintained in normal culture conditions up until the point of video recording. Individual larva were picked and placed onto the centre of an agar plate at room temperature and recorded immediately. Video plate recordings were carried out with use of a Teledyne DALSA Genie Nano Camera (G3-GM11-M2420) using Gecko GigE Video Recorder software (v2.0.3.1; www.visionexperts.co.uk), at 25 frames per second, ensuring a sharp contrast between the larva and background. 20 individual larvae were recorded per condition up to a maximum of 5 minutes or until the larva was no longer in the field of view (FOV). Individual videos–each corresponding to one larva–were segmented, tracked and skeletonised using Tierpsy Tracker^12^. Data were discarded from subsequent analysis if video tracking proved unsuccesssful, which occurred when the head was incorrectly identified in 12 videos. When the head was correctly identified it was stable over the course of the video. 256 defined features, previously determined to be useful in classifying behaviour in *C.elegans*, were extracted from the tracking data and compared between conditions^13^. Correction for multiple testing was applied using the Benjamini–Hochberg procedure to control the false discovery rate to 0.05^31^.

### Quantitative RT-PCR

Larvae were washed in PBS and dissected in RNALater solution. Total RNA was extracted from pooled dissected tissue using the QIAGEN RNeasy Mini Kit. First-strand DNA was synthesised with the iScript cDNA synthesis kit (Bio-Rad). PCR was performed by mixing cDNA samples with iTaq Universal SYBR Green Supermix (Bio-Rad), ROX passive reference dye (Bio-Rad) and the relevant primers in a 96-well plate. Analysis was carried out on a 7900HT real-time PCR system (Applied Biosystems). At least 3 independent biological replicates and 2 technical replicates were used. Expression values were normalised to *RpL32* or *Act88F*. Primers used are listed in Extended Data Table 1.

### RNA-Sequencing

RNA from pooled dissected tissue was extracted with the QIAGEN RNeasy Mini Kit. A minimum of 3 samples were prepared for each genotype. RNA-seq libraries were prepared from a minimum of 10ng of total RNA using the Illumina Truseq mRNA stranded library prep kit (Illumina, San Diego, USA) according to the manufacturer’s protocol. Library quality was checked on a Bioanalyser HS DNA chip and concentrations were estimated by Qubit measurement. Libraries were pooled in equimolar quantities and sequenced on a Hiseq2500 using paired end 100bp reads. At least 35 million reads passing filter were achieved per sample. After demultiplexing, raw or trimmed RNA-Seq reads were aligned against Ensembl *Drosophila* genome reference sequence assembly (dm3) and transcript annotations using TopHat2^32^. Trim Galore, developed at The Babraham Institute by Felix Krueger (http://www.bioinformatics.babraham.ac.uk/projects/trim_galore/), was applied for trimming adapter and low quality reads. For differential gene expression analysis, gene-based read counts were then obtained using the featureCounts function from Rsubread Bioconductor package^33^. Differential expression analysis was performed on the counts data using DESeq2 Bioconductor package^34^. From tumour RNA-sequencing data, putative tumour-secreted factors were annotated based on identification of genes labelled with the GO term ‘extracellular’ or presence of the signal peptide sequence as identified by the signal peptide database (http://www.signalpeptide.de).

### Gene Set Enrichment Analysis

GSEA was carried out using a ranked gene list based on Wald statistics from DESeq2 results^35^. MSigDB gene sets from ‘H’, ‘C2’ and ‘C5’ collections were used and only considered where FDR q value <0.25. Normalised enrichment scores were calculated for multiple terms within functional categories. Results were displayed as cumulative normalised enrichment score, as previously published^36^.

### Hemolymph sample preparation for metabolomic analyses

Pooled hemolymph was collected from multiple animals on ice to a combined minimum sample volume of 25 µl. Each sample was mixed with 225 µL of methanol containing internal standards (50 µM). Then, chloroform (250 µL) and Milli-Q water (100 µL) were added, mixed and centrifuged. The water layer was filtrated through a 5-kDa cut-off filter to remove macromolecules. Filtrate was centrifugally concentrated and resuspended in 25 µL of ultrapure water prior to measurement. n=4 per condition.

### Metabolomic analyses

Targeted metabolomic analyses were performed by the Human Metabolome Technologies Inc., Yamagata, Japan. Capillary electrophoresis mass spectrometry (CE-MS) was performed using Agilent CE-TOFMS Machine (CE-TOFMS) and Agilent 6460 TripleQuad LC/MS Machine (CE-QqQMS)(Agilent Technologies). Compounds were measured based on previously described methods^37-39^. Peaks detected in CE-TOFMS analysis were extracted using automatic integration software (MasterHands ver.2.17.1.11 developed at Keio University)^40^ and those in CE-QqQMS analysis were extracted using automatic integration software (MassHunter Quantitative Analysis B.06.00 Agilent Technologies, Santa Clara, CA, USA) in order to obtain peak information including m/z, migration time (MT), and peak area. Putative metabolites were assigned from peak alignments based on the HMT metabolite database on the basis of m/z and MT. Relative peak values were calculated based on internal standards and normalised to sample volumes. Absolute quantification was calculated by normalising the peak area of each metabolite with respect to the area of the internal standard and by using standard curves, which were obtained by three-point calibrations. For hemolymph samples, 214 metabolites (135 metabolites in cation mode and 79 metabolites in anion mode respectively) were annotated based on the HMT metabolite database, including 3-methylhistidine and proline.

### Human cancer database analyses

TCGA IlluminaHiSeq_RNASeqV2 datasets for Colon Adenocarcinoma (COAD), Stomach Adenocarcinoma (STAD), Pancreatic Adenocarcinoma (PAAD), Rectum Adenocarcinoma (READ), Liver Hepatocellular Carcinoma (LIHC), Kidney Renal Clear Cell Carcinoma (KIRC), Kidney Chromophobe (KICH) and Kidney Renal Papillary Cell Carcinoma (KIRP) were obtained via the GDC Legacy Archive. Corresponding patient clinical data, including overall survival status was obtained from cBioPortal for Cancer Genomics (http://www.cbioportal.org.)^41,42^. When combining overlapping data from both datasets the following number of patients remained and were analysed with respect to each cancer type: (COAD – 177; STAD - 261; PAAD – 179; READ – 66; LIHC – 373; KIRC – 534; KICH – 66 and KIRP - 291). The three kidney cancer tumour types (KIRC, KICH and KIRP) were combined into a pooled kidney cancer group with a total of 891 patients.

Gene expression values were taken from the RSEM RNA-Sequencing data. The R/Bioconductor package edgeR (3.22.3) was applied for trimmed mean of M-values (TMM) normalisation with the calcNormFactors function. The function voom from R/Bioconductor package limma (3.36.3) was used for converting raw counts to a log2 scale (counts per million). Individual patients were assigned a Cachexia Score based on median expression levels of the following cachectic factors: IL-1A, IL-1B, IL-6, IL-8 and TNF-α. Pearson correlation coefficients were calculated for six cancer types between the Cachexia Score and expression of either SLC36A1 or SLC36A4. The SLC36A2 and SLC36A3 isoforms of the SLC36A gene were excluded from analysis due to their low expression levels across the tumour types examined. The Benjamini & Hochberg adjusted p-value was calculated to measure the statistical significance of observed correlations as denoted by the False Discovery Rate (FDR)^31^.

For cancer types where statistically significant positive correlation was observed (FDR<0.05), we defined two patient groups—(i) with high Cachexia Score and high SLC36 transporter expression (High; red) and (ii) with low Cachexia Score and low SLC36 transporter expression (Low; blue). Each group represents the upper (High) and lower (Low) quartile of patients in each cancer type, as calculated from linear regression analysis using SLC36 expression level as the explanatory variable. Kaplan-Meier survival curves were used to compare survival outcomes between the High and Low patient groups within each cancer type^43^. The Log Rank test was used to assess the statistical significance of survival curves and calculated p-values are displayed.

### Statistical methods

Unless otherwise stated, all statistical analyses were carried out with an unpaired, two-tailed students t-test in GraphPad Prism software (v.7.0d) where p<0.005 = ***; 0.005<p<0.025 = **; 0.025<p<0.05 = *.

## ACKNOWLEDGEMENTS

We thank members of the Hirabayashi, Miguel-Aliaga, and Cochemé laboratories for helpful discussions; members of the Genomics facility (L. Game, K. Rekopoulou, and I. Andrew) for help with RNA-sequencing; C.Choutka, J. Cordero and I. Miguel-Aliaga for manuscript comments; Luigi Feriani for technical help with larval video tracking and analysis, and A. O’Reilly, M. Simon, G. Halder, M. Krasnow, H. Stocker, B. Shilo, T. Koyama, J. Parrish, and D. Goberdhan for kindly providing reagents. We also thank the Bloomington Drosophila Stock Center (NIH P40OD018537), and Vienna Drosophila Resource Center (VDRC, www.vdrc.at) for fly strains. A.E.X.B. was supported intramural funding from the Medical Research Council (MC-A658-5TY30). S.H. was supported by grants from PRESTO, Japan Science and Technology Agency and intramural funding from the Medical Research Council (MC-A654-5QC30).

## AUTHOR CONTRIBUTIONS

H.N., L.C., A.E.X.B. and S.H. performed and analysed experiments. Y-F. W. carried out bioinformatics analysis. H.N. and S.H. conceived and designed the project and wrote the manuscript, with all authors providing feedback.

---

**Extended Data Figure 1.**
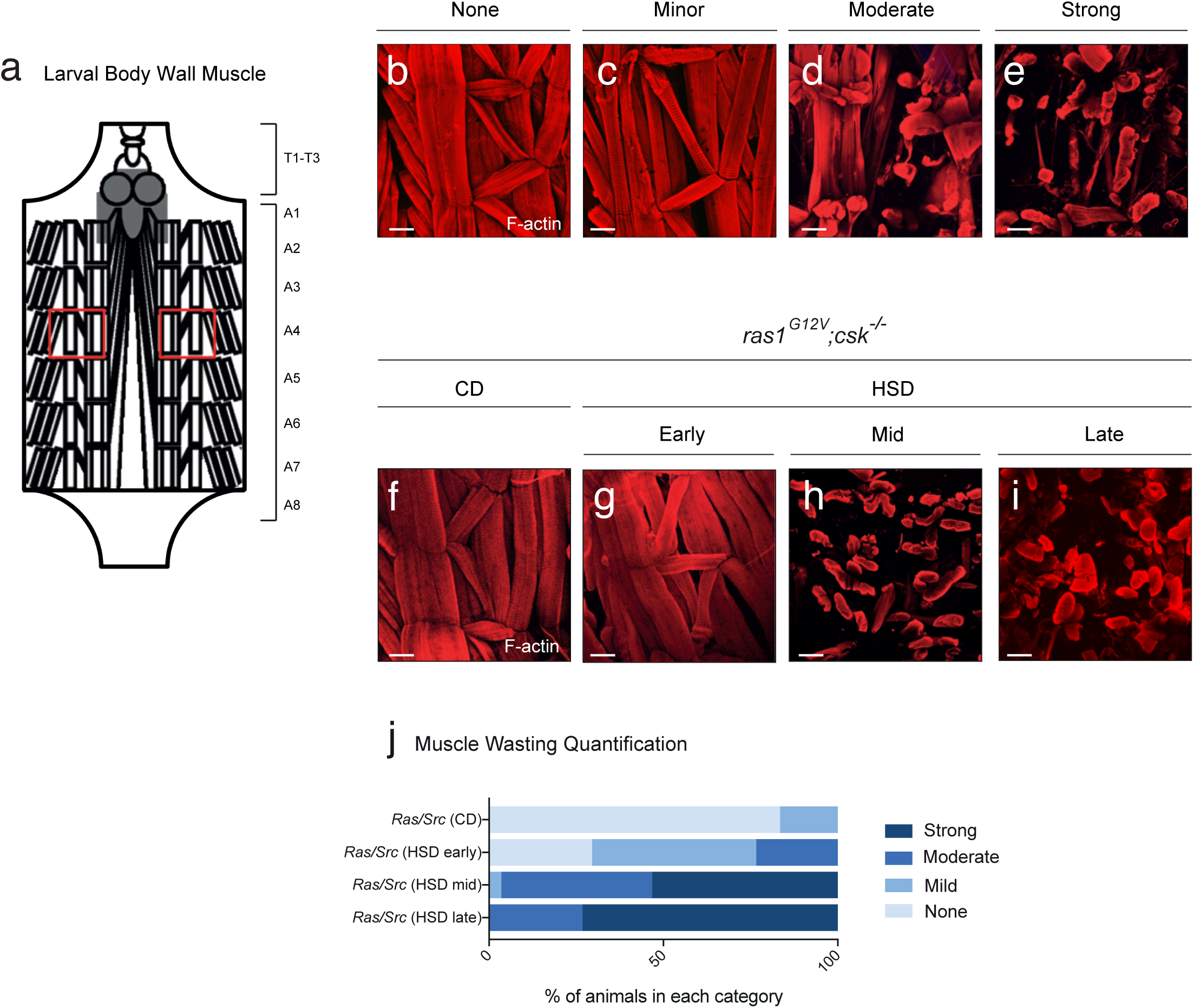
HSD-fed Ras/Src-animals exhibit progressive cachexia-like muscle wasting. **a**, Schematic of larval body wall muscle upon dissection from anterior (top) to posterior (bottom). The larval body wall contains 3 thoracic segments (T1-T3) and 9 abdominal segments (A1-A9). Abdominal muscles from segments A1-A8 are labelled. Red boxes outline regions of the body wall muscle which are imaged, containing the ventral longitudinal, lateral oblique and lateral longitudinal muscles centred around the fourth abdominal segment. **b-e**, Representative images of dissected larval body wall muscle stained with phalloidin to visualise F-actin. Muscle wasting scores increase in strength from left to right: “no wasting” (**b**), “minor” (**c**), “moderate” (**d**) and “strong” (**e**). Scale bar, 100 μm. **f-i**, Dissected larval body wall muscle of *ras1*^*G12V*^;*csk*^*-/-*^animals raised on CD (**f**) or on HSD at Early (**g**), Mid (**h**) and Late stage (**i**) Scale bar, 100 μm. **j**, Matching body wall muscle wasting quantification of *ras1*^*G12V*^;*csk*^*-/-*^ animals raised on CD and HSD. X-axis represents the percentage of individual animals scoring in each category denoted “none”, “mild”, “moderate” or “strong”. n≥30 per condition.

**Extended Data Figure 2.**
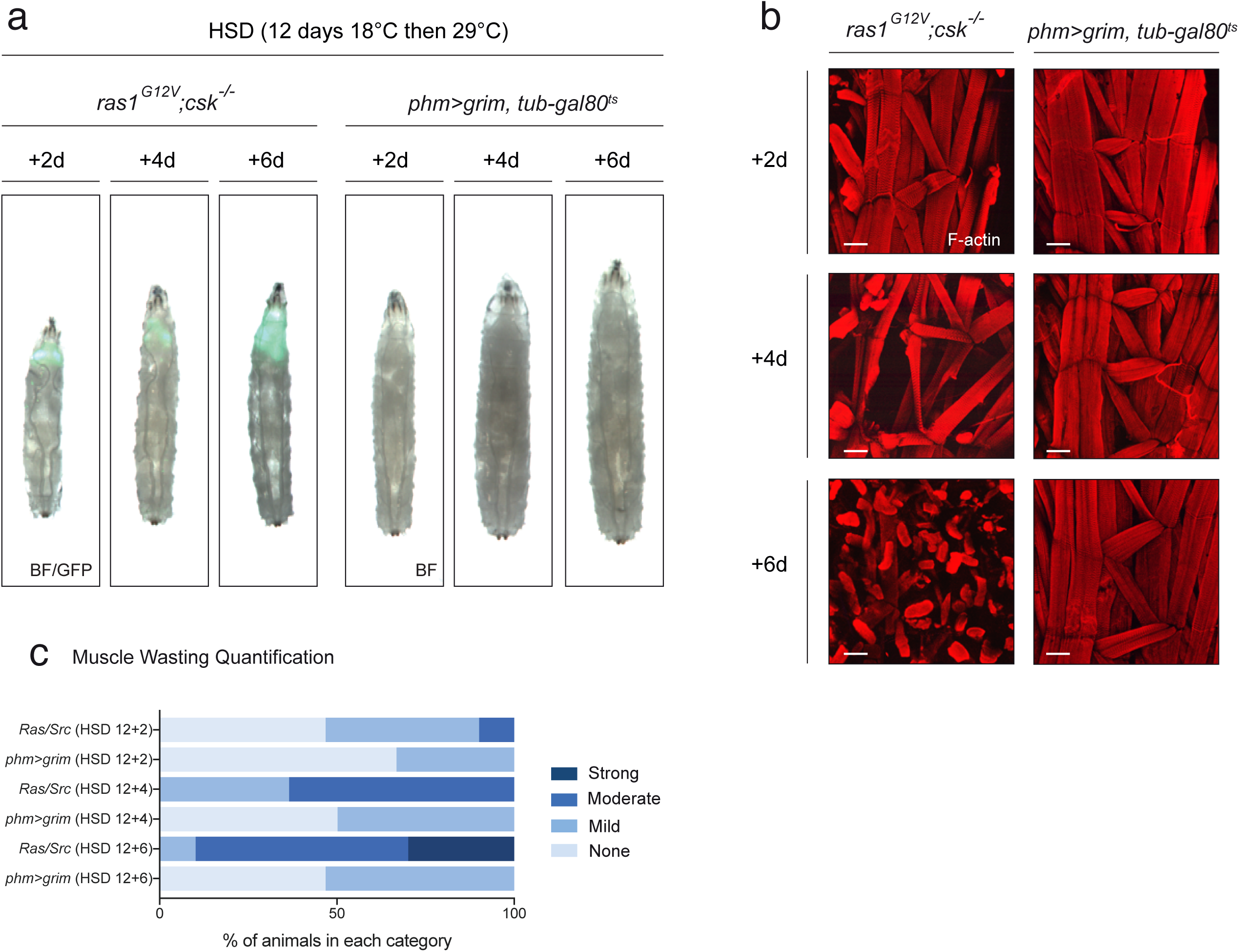
Extended larval period is insufficient to account for systemic muscle wasting. **a**, Third instar larvae of *ras1*^*G12V*^;*csk*^*-/-*^ and *phm>grim, tub-gal80*^*ts*^ animals raised on HSD for 12 days in 18°C plus 2, 4 and 6 days after temperature shift to 29°C (+2d, +4d and +6d respectively). *ras1*^*G12V*^;*csk*^*-/-*^ animals are imaged under BF and GFP. *phm>grim* animals (tumour-free) are imaged under BF. **b**, Matching dissected larval body wall muscle. Scale bar, 100 μm. **c**, Matching body wall muscle wasting quantification. n≥30 per condition.

**Extended Data Figure 3.**
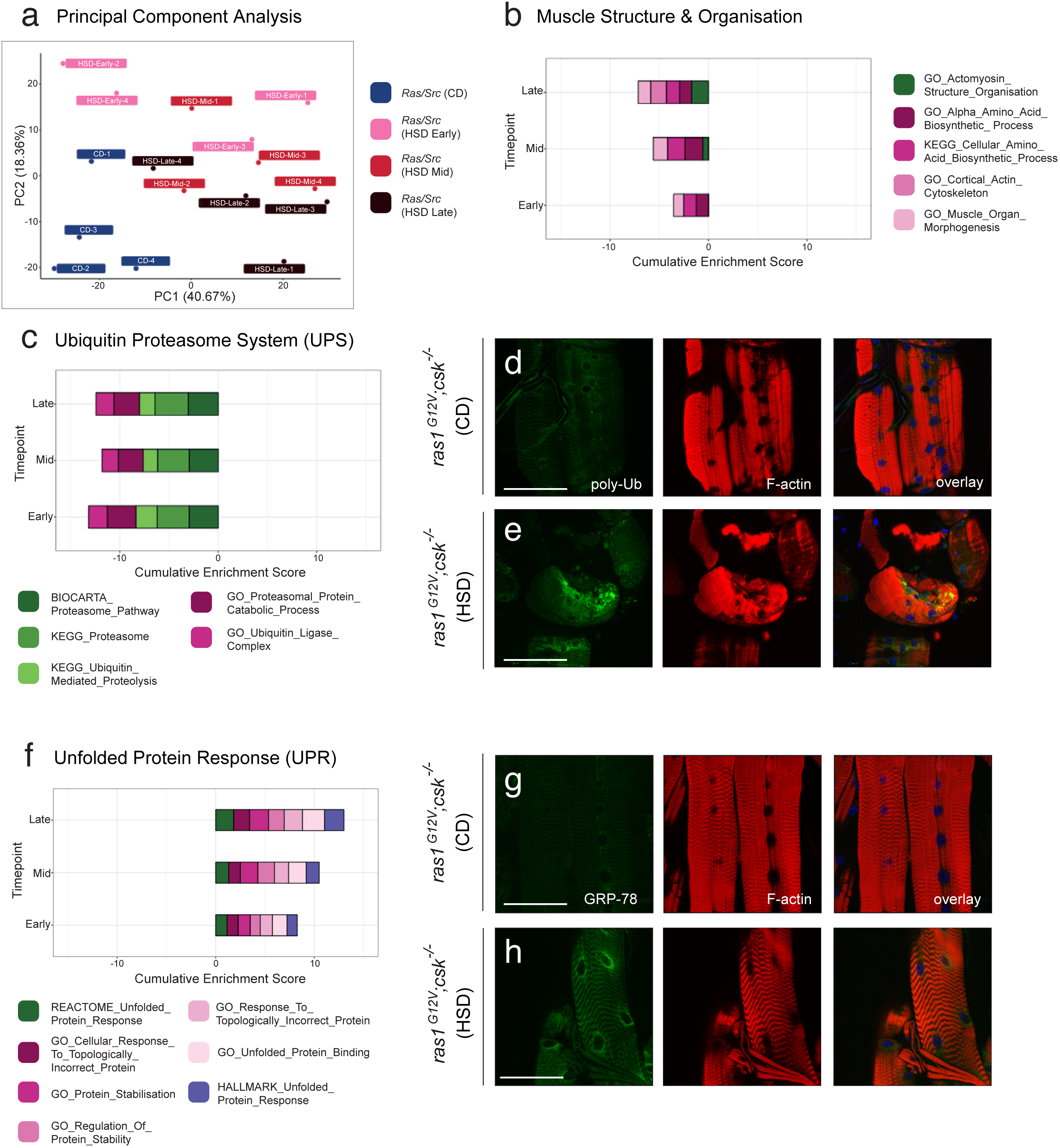
HSD-fed Ras/Src-animals promote muscle wasting via proteasome dysfunction and unfolded protein response. **a**, Principal Component Analysis of dissected muscle RNA-sequencing data from *ras1*^*G12V*^;*csk*^*-/-*^ animals raised on CD (blue) and *ras1*^*G12V*^;*csk*^*-/-*^ animals raised on HSD at Early (pink), Mid (red) and Late Stage (brown). n=4 per condition. **b**, Gene Set Enrichment Analysis (GSEA) for the functional category ‘Muscle Structure & Organisation’ from dissected muscle RNA-sequencing data from *ras1*^*G12V*^;*csk*^*-/-*^ animals raised on HSD at Early Stage (Early), Mid Stage (Mid) and Late Stage (Late) compared to *ras1*^*G12V*^;*csk*^*-/-*^ animals raised on CD. Data is displayed as cumulative enrichment score over the time course. Gene sets in purple are derived from Hallmark (H) datasets, gene sets in green are derived from Curated (C2) datasets and gene sets in pink are derived from Gene Ontology (C5) datasets. **c**, GSEA (as for **b**) for the functional category ‘Ubiquitin Proteasome System’. **d, e**, Poly-Ubiquitin staining (green) of dissected body wall muscle from *ras1*^*G12V*^;*csk*^*-/-*^ animals raised on CD (**d**) or HSD (**e**) and with DAPI (blue) and F-actin (red). Scale bar, 100 μm. **f**, GSEA (as for **b**) for the functional category ‘Unfolded Protein Response’. **g, h**, GRP-78 staining (green) of dissected body wall muscle from *ras1*^*G12V*^;*csk*^*-/-*^ animals raised on CD (**g**) or HSD (**h**) and with DAPI (blue) and F-actin (red). Scale bar, 100 μm.

**Extended Data Figure 4.**
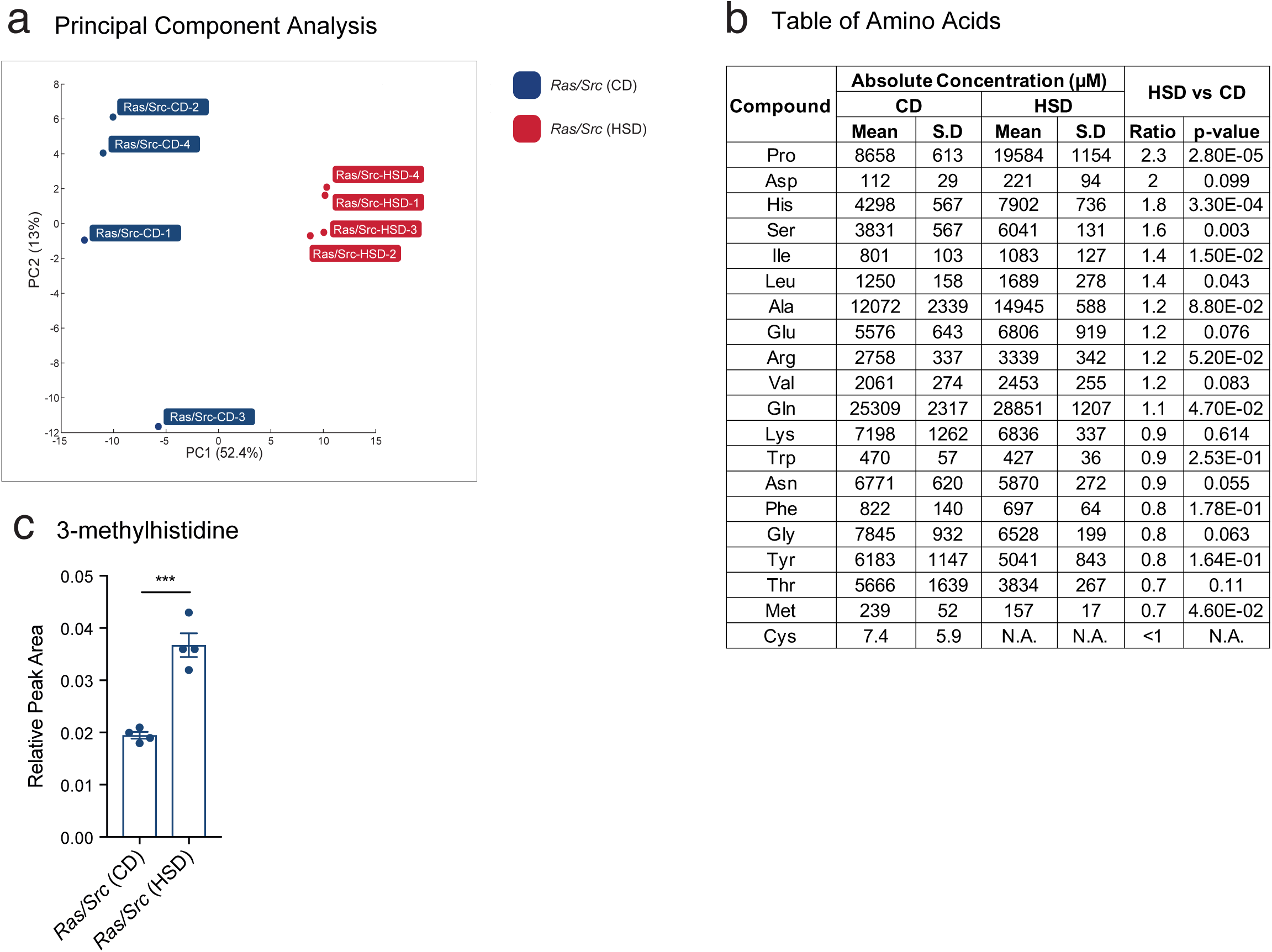
Metabolomic analysis of hemolymph reveal an increase in circulating amino acids in HSD-fed Ras/Src-animals. **a**, Principal Component Analysis of targeted metabolomic analysis from hemolymph in *ras1*^*G12V*^;*csk*^*-/-*^animals raised on CD (blue) or HSD (red). n=4 per condition. **b**, Amino acid concentrations from targeted metabolomic analysis of hemolymph in *ras1*^*G12V*^;*csk*^*-/-*^ animals raised on CD (blue) or HSD (red). n=4 per condition. Amino acids are listed in order of highest fold-elevation in *ras1*^*G12V*^;*csk*^*-/-*^ animals raised on HSD compared to CD. **c**, Circulating 3-methylhistidine levels depicted by relative peak area from hemolymph in *ras1*^*G12V*^;*csk*^*-/-*^animals raised on CD or HSD. Results are shown as mean ± SEM. Asterisks indicate statistically significant difference (***; p<0.005).

**Extended Data Figure 5.**
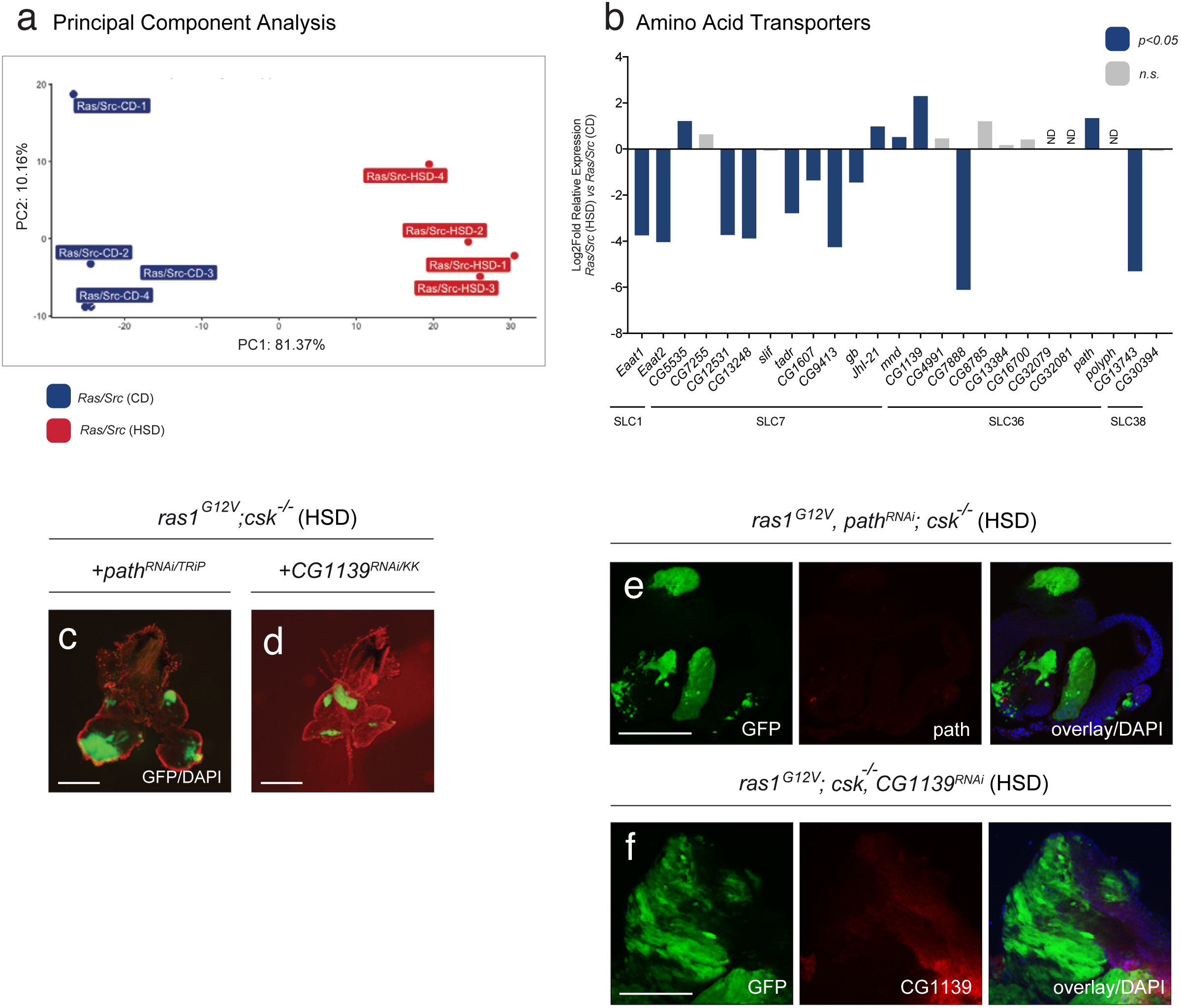
RNA-Seqencing identifies candidate amino acid transporters required for diet-enhanced tumour growth. **a**, Principal Component Analysis of RNA-sequencing data from dissected tumour tissue in *ras1*^*G12V*^;*csk*^*-/-*^ animals raised on CD (blue) or HSD (red). n=4 per condition. **b**, Expression levels of amino acid transporters of the SLC1, SLC7, SLC36, SLC38 families as measured by RNA-seq analysis. Gene expression levels are shown as the log2-relative fold change between *ras1*^*G12V*^;*csk*^*-/-*^ animals raised on HSD compared to *ras1*^*G12V*^;*csk*^*-/-*^ animals raised on CD. Blue bars indicate cases where p<0.05. Grey bars indicate cases where p>0.05. **c, d**, Dissected eye epithelial tissue stained with DAPI (red) from animals raised on HSD with independent RNAi lines, *ras1*^*G12V*^, *path*^*RNAi/TRiP*^;*csk*^*-/-*^ (**c**) or *ras1*^*G12V*^,*CG1139*^*RNAi/KK*^;*csk*^*-/-*^ (**d**). Scale bar, 250 μm. **e**, Anti-Path staining (red) of dissected tumour tissue from *ras1*^*G12V*^, *path*^*RNAi/KK*^;*csk*^*-/-*^animals raised on HSD with DAPI (blue). Scale bar, 100 μm. **f**, Anti-CG1139 staining (red) of dissected tumour tissue from *ras1*^*G12V*^;*csk*^*-/-*^,*CG1139*^*RNAi/GD*^ animals raised on HSD, with DAPI (blue). Scale bar, 100 μm.

**Extended Data Figure 6.**
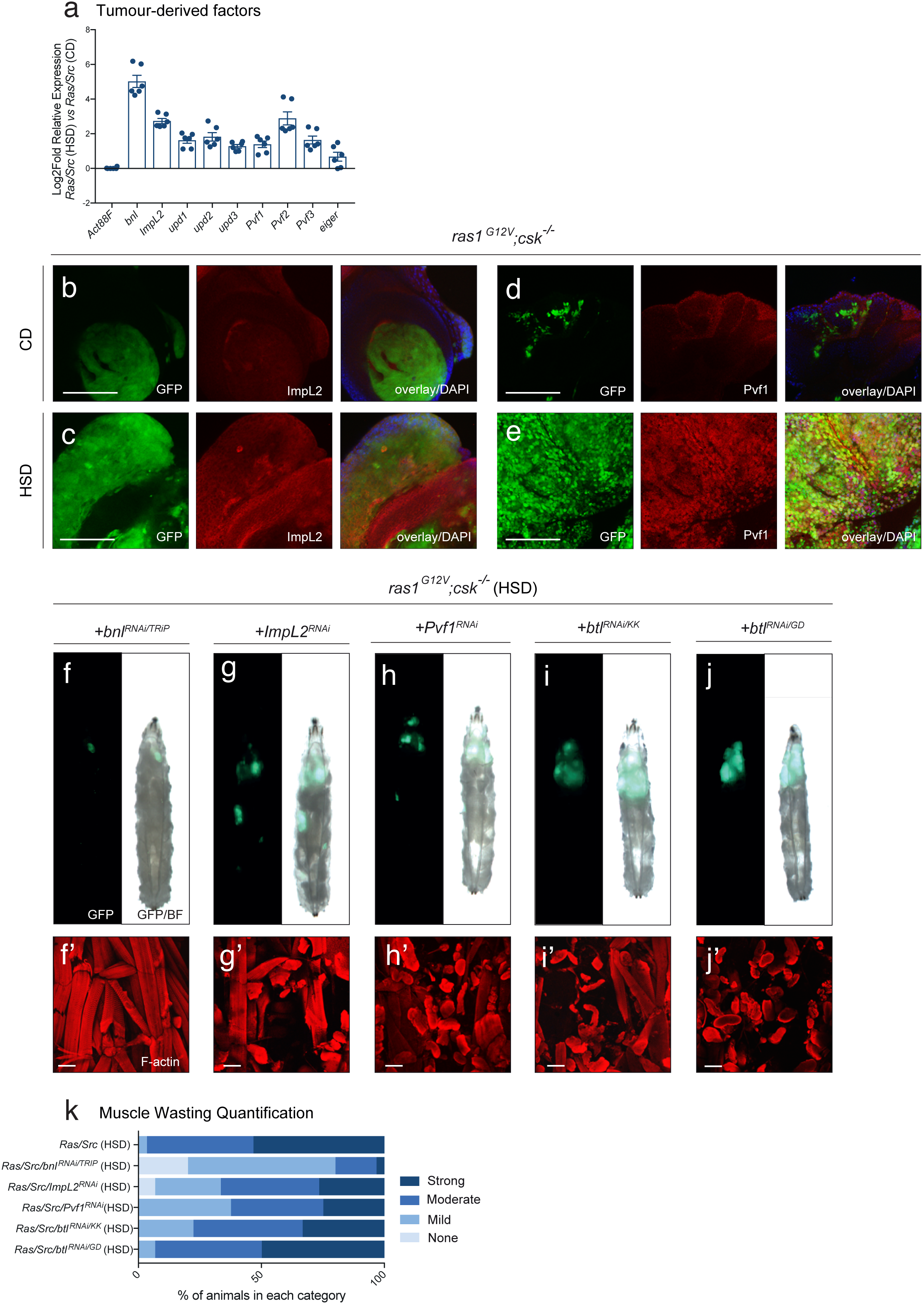
RNA-Seqencing identifies branchless as a tumour-derived factor that contributes to muscle wasting and tumour growth. **a**, qPCR validation of candidate tumour-derived factors—*bnl, ImpL2, upd1, upd2, upd3, Pvf1, Pvf2, Pvf3* and *eiger*—initially identified by RNA-sequencing. Gene expression levels are shown as log2-relative fold change between *ras1*^*G12V*^;*csk*^*-/-*^ animals raised on HSD compared to *ras1*^*G12V*^;*csk*^*-/-*^ animals raised on CD, as determined by qPCR. Samples are normalised to *Act88F*. Results are shown as mean ± SEM. **b, c**, ImpL2 staining (red) of dissected tumour tissue from *ras1*^*G12V*^;*csk*^*-/-*^ animals raised on CD (**b**) or HSD (**c**) with DAPI (blue). Scale bar, 100 μm. **d, e**, Pvf1 staining (red) of dissected tumour tissue from *ras1*^*G12V*^;*csk*^*-/-*^ animals raised on CD (**d**) or HSD (**e**) with DAPI (blue). Scale bar, 100 μm. **f-j**, *ras1*^*G12V*^;*csk*^*-/-*^,*bnl*^*RNAi/TRIP*^ (**f**), *ras1*^*G12V*^;*csk*^*-/-*^,*ImpL2*^*RNAi*^ (**g**), *ras1*^*G12V*^;*csk*^*-/-*^,*Pvf1*^*RNAi*^ (**h**), *ras1*^*G12V*^;*csk*^*-/-*^,*btl*^*RNAi/KK*^ (**i**), and *ras1*^*G12V*^;*csk*^*-/-*^,*btl*^*RNAi/GD*^ (**j**) third-instar larvae raised on HSD. **f’-j’**, Matching dissected larval body wall muscle. Scale bar, 100 μm. **k**, Matching body wall muscle wasting quantification. n≥30 per condition.

**Extended Data Figure 7.**
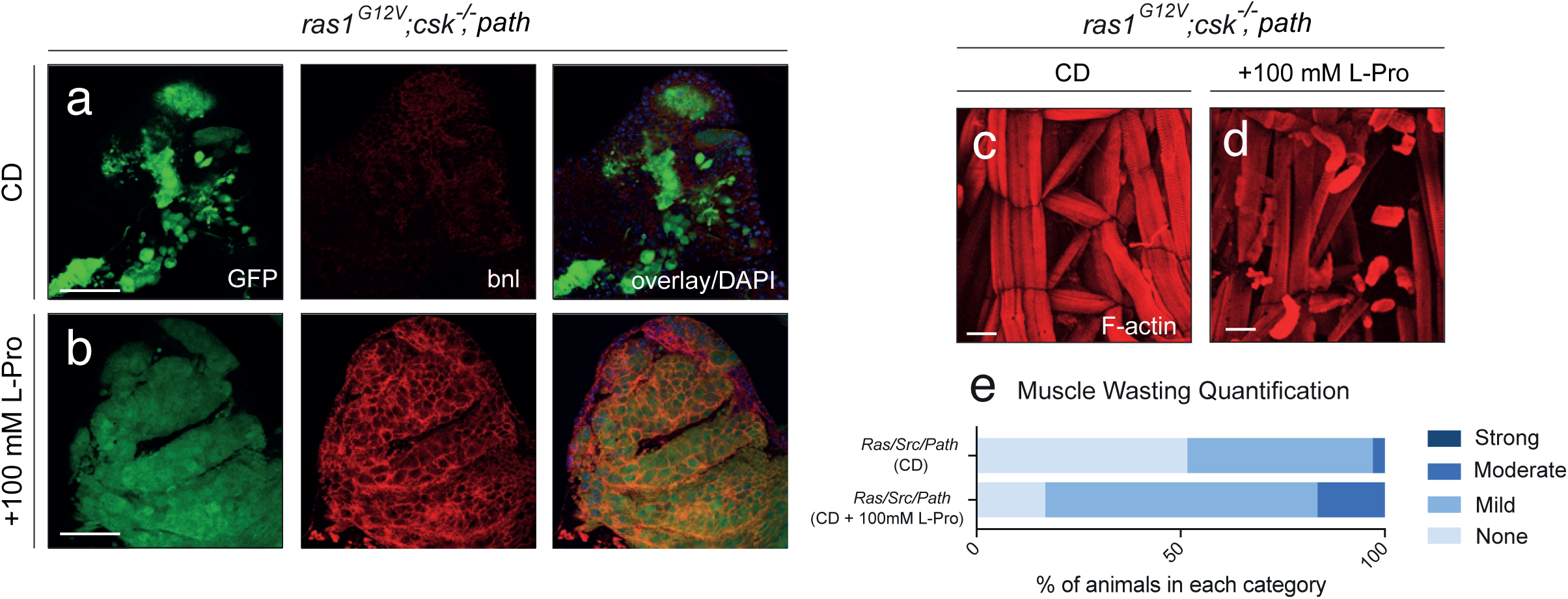
Proline-mediated Ras/Src tumour growth promotes Bnl-expression and muscle wasting. **a, b**, Anti-Bnl staining (red) of dissected tumour tissue from *ras1*^*G12V*^;*csk*^*-/-*^,*PathA* animals raised on CD (**a**) or CD supplemented with 100mM proline (**b**) with DAPI (blue). Scale bar, 40 μm. **c, d**, Dissected larval body wall muscle from *ras1*^*G12V*^;*csk*^*-/-*^,*PathA* animals raised on CD (**c**) or CD supplemented with 100mM proline (**d**). **e**, Matching body wall muscle wasting quantification. n≥30 per condition.

**Extended Data Table 1.**
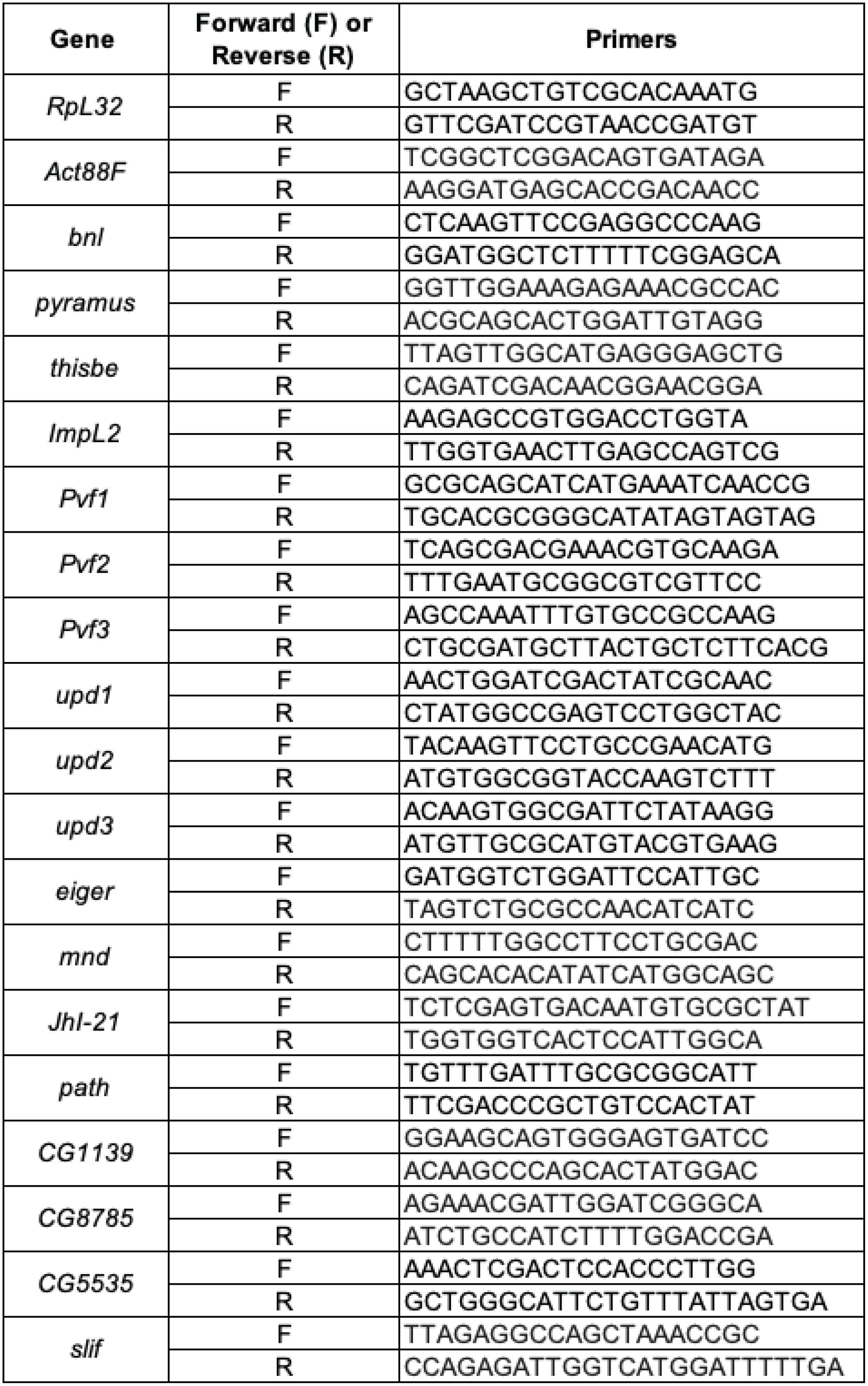
List of primers used in this study.

